# Machine learning enables *de novo* multi-epitope design of plasmodium falciparum circumsporozoite protein to target trimeric L9 antibody

**DOI:** 10.1101/2025.06.29.662177

**Authors:** J. Andrew D. Nelson, Samuel E. Garfinkle, Zi Jie Lin, Joyce Park, Amber J. Kim, Kelly Bayruns, Madison E. McCanna, Kylie M. Konrath, Colby J. Agostino, Daniel W. Kulp, Audrey R. Odom John, Jesper Pallesen

## Abstract

Currently approved vaccines for the prevention of malaria provide only partial protection against disease, due to high variability in the quality of induced antibodies. These vaccines present the unstructured central repeat region, as well as the C-terminal domain, of the circumsporozoite protein (*Pf*CSP) of the malaria parasite, *Plasmodium falciparum* (1). A recently discovered protective monoclonal antibody, L9, recognizes three structured copies of the *Pf*CSP minor repeat. Similarly to other highly potent anti-malarial antibodies, L9 relies on critical homotypic interactions between antibodies for its high protective efficacy (2, 3). Here, we report the design of immunogens scaffolding one copy of *Pf*CSP’s minor repeat capable of binding L9. To design immunogens capable of presenting multiple, structure-based epitopes in one scaffold, we developed a machine-learning-driven structural immunogen design pipeline, MESODID, tailored to focus on multi-epitope vaccine targets. We use this pipeline to design multiple scaffolds that present three copies of the *Pf*CSP minor repeat. A 3.6 Å cryo-EM structure of our top design, minor repeat targeting immunogen (M-TIM), demonstrates that M-TIM successfully orients three copies of L9, effectively recapitulating its critical homotypic interactions. The wide prevalence of repeated epitopes in key vaccine targets, such as HIV-1 Envelope, SARS-CoV-2 spike, and Influenza Hemagglutinin, suggests that MESODID will have broad utility in creating immunogens that incorporate such epitopes, offering a new powerful approach to developing vaccines against a range of challenging infections, including malaria.

**Significance Statement:** In this study, we present a machine learning driven, structure-guided protein design pipeline built specifically for the design of multi-epitope vaccine immunogens. We employ the pipeline here to design and solve the cryo-EM structure of a *de novo* immunogen that binds and properly orients three copies of the anti-malarial monoclonal antibody L9, producing a promising next-generation malaria vaccine immunogen. This design pipeline could be employed to design any number of structurally constrained multi-epitope immunogens, as well as other proteins designed to bind multiple targets.

## Introduction

Malaria is an infectious disease caused by apicomplexan parasites of the genus *Plasmodium* that continues to kill 600,000 people annually (4). *Plasmodium falciparum (Pf)* is especially deadly and is responsible for the majority of global malaria mortality (4). While much progress has been made in the past 25 years, resistance to first-line insecticides (5) and anti-malarial drugs (6) threaten the advances that have been made in malaria control. Vaccines are a valuable tool that can provide much-needed help in the fight against malaria. Recent years have seen the approval of the world’s first two vaccines for malaria by the World Health Organization (WHO), as well as their implementation within multiple countries’ vaccination strategies (7, 8).

The lifecycle of the *Pf* parasite begins when sporozoites are transmitted through a mosquito bite and travel to the liver, where they mature and initiate the pre-erythrocytic stage of infection (9). Both malaria vaccines approved thus far, RTS,S/AS01 (RTS,S) and R21/MATRIX-M (R21), target the pre-erythrocytic stage of the parasite, and specifically the circumsporozoite protein (CSP) (10, 11). CSP is the most ubiquitous protein on the surface of sporozoites and broadly consists of 3 domains: an N-terminal domain critical for recognizing heparan sulfate proteoglycans in the liver; an immunodominant central repeat region consisting of repetitive four-amino-acid motifs; and a C-terminal thrombospondin-like adhesive domain (12–14). The central repeat region consists of both major (NANP/NPNA) and minor (NVDP/NPNV) repeats, so called because of the relative abundance of each repeat within the central repeat region (15). RTS,S and R21 both consist of 19 copies of the major repeat as well as the C-terminus of CSP fused to the Hepatitis B Surface Antigen (HbsAg), forming virus like particles (VLPs) that present disordered NANP/NPNA-based motifs when expressed in yeast. RTS,S and R21 differ only in the frequency of CSP-based epitopes presented on the surface of VLPs and in the adjuvants used (16). Both vaccines exhibit suboptimal efficacy due to variability in the potency of elicited antibodies (1). Thus, these landmark vaccines could still be further optimized or supplemented to improve both the quality and durability of antibody responses over time.

To interrogate the antibody responses which are most effective in preventing malaria transmission, groups have isolated and structurally characterized a number of monoclonal antibodies that target CSP (1, 17–20)’. Of interest, the human heavy chain germline IGHV3-33 has been implicated in the majority of the most potent anti-CSP antibodies isolated thus far (21). These IGHV3-33-derived antibodies often develop inter-fab homotypic interactions between antibodies, which are critical for the antibodies’ protective efficacy and are selected for via somatic hypermutation within germinal centers (22, 23)’. One of the most potent monoclonal antibodies isolated thus far, L9, is derived from IGHV3-33 and relies on homotypic interactions for its protective efficacy *in vivo,* specifically targeting the minor (NPNV) repeat (2, 18). Antibodies with the same potency and specificity as L9 likely remain rare in the polyclonal antibody response to vaccination given the absence of the minor repeat in the approved vaccines, the high titers of anti-CSP antibodies required for protection against disease (16), and the low frequency of somatic hypermutation seen in antibodies induced by vaccination (24). The central repeat region of CSP is highly flexible and disordered (12, 25)’, but can adopt stable, structured conformations upon antibody binding (26). If portions of the central repeat region of CSP could be pre-configured in a structurally stabilized conformation capable of binding germline precursors of homotypically interacting antibodies, we hypothesize that this could result in the induction of more potent structure-specific, homotypically interacting anti-CSP antibodies, such as L9 (2). However, vaccine design tools focused on developing antigens able to arrange antibodies that mimic these homotypic interactions have not been available.

Recent years have also seen rapid advances in the application of machine learning to protein design, highlighted by the development of tools such as RFdiffusion (27) and ProteinMPNN (28) for protein backbone generation and sequence design, respectively. These tools enable design of protein scaffolds of diverse, complicated topologies with unprecedented accuracy. However, these tools can require additional refinement of designs beyond their initial output, and scaffolding repetitive, structure-based epitopes provides added difficulty due to structural constraints. To address these limitations, we developed MESODID (Multi-Epitope Structure-Optimized Deep Iterative Design), an iterative design pipeline tailored specifically to refine machine-learning-based protein designs that scaffold repeat epitopes. We use this pipeline to scaffold three discontinuous minor repeat motifs in the same relative orientation observed in cryo-EM structures (2, 3)’ of CSP in complex with three copies of monoclonal L9. We show that scaffolds designed using this methodology express and bind L9. Our top design, minor repeat targeting immunogen (M-TIM), binds three copies of L9 in the same orientation observed in prior cryo-EM structures with CSP, effectively recapitulating the homotypic interactions of the wild type CSP-L9 complex.

## Results

### DESIGN, EXPRESSION, AND ANTIGENIC PROFILING OF SINGLE MINOR REPEAT IMMUNOGENS

To examine if the standard AI design pipeline could scaffold minor-repeat-containing motifs, we first designed immunogens that scaffold a single minor repeat motif using RFdiffusion (27) and ProteinMPNN (28) with ESMFold (29) as a structure prediction oracle. A total of 25 designs (Table S1) were manually selected based on high predicted local distance difference test (pLDDT) and low alpha carbon root mean square deviation (C_α_-RMSD) to the minor repeat epitope from a recently published structure of the L9-CSP complex (PDB 8EK1) (2). In particular, all of the single repeat immunogens selected had a pLDDT > 85 and epitope RMSD < 1 Å (average pLDDT of 85.9, RMSD of 0.2 Å). Of the 25 designs ordered, 11 expressed with yields of 0.1-1.8 mg protein/100 mL transfection culture, and 9 of these designs could be isolated by SEC (Figure S1A). We next assessed binding to L9 by ELISA, and we observed that all 9 designs isolated by SEC bound to L9, with no binding seen to an HIV-1 Envelope trimer, an unrelated negative control (Figure S1B). This confirms that standard deep learning protein design methods can be deployed to design minor-repeat-based scaffolds. Encouraged by these results, we applied the same design process to generate scaffolds for triple minor repeat immunogens. Reflecting the increased challenge of scaffolding multiple epitopes in a fixed 3D conformation, none of the initial round of triple repeat designs featured pLDDT > 85 and epitope RMSD < 1 Å (average pLDDT of 81.2, RMSD of 1.28 Å). Given the modest success rate from the first round of designs, we were concerned that our initial triple minor repeat scaffolds would not reliably express and fold into a 3D conformation capable of recapitulating homotypic interactions with L9. To further refine our designs of the three-copy minor repeat conformational epitope, we developed the MESODID pipeline for design of multi-epitope immunogens.

### MESODID PIPELINE

The MESODID pipeline (Figure 1A,1B) was built with the concept of iterating deep learning design with structure prediction. The pipeline begins with deep-learning based backbone (RFdiffusion (27)) and sequence (ProteinMPNN (28)) generation, followed by deep-learning structure prediction (ESMFold (29)). We built three refinement modules to improve these iterative, predicted design intermediates (saturation mutagenesis, disulfide engineering, and energy filtering). The saturation mutagenesis module mutates every amino acid outside the epitope of interest to every other possible amino acid; we found that single mutations can provide improvements in predicted local distance difference test (pLDDT) (Figure 1C). We found additional pLDDT improvements through the disulfide engineering module, which uses the Molecular Software Library (MSL) (30) to efficiently identify residue pairs within the designed scaffold that are structurally homologous to a large dataset of naturally occurring disulfide bonds derived from the Protein Data Bank (Figure 1D). Designs are filtered by a weighted score that incorporates contributions from both pLDDT and the RMSD of the selected epitope to its coordinates in its reference structure. We found that 24.1% of saturation mutagenesis single mutants and 11.6% of disulfide mutants improved the predicted weighted score (Figure S2A-B). Iterative rounds of refinement where top mutations are recombined were found to be key to improving pLDDT/RMSD metrics (Figure 1E). Designs are then passed through a module that sorts according to energetic favorability. In the energy module, individual designs are aligned to the structural epitope within the reference structure (PDB 8EK1) and the entire complex (candidate immunogen design + bound antibodies) undergoes constrained relaxation in Rosetta (31–34) to abrogate potential detrimental clashes between the immunogen and antibodies of interest that may prevent binding (Figure 1F). Rosetta Score post-relaxation of the entire complex is used as a further filter on top designs. This iterative refinement continues for multiple rounds (here we used up to six), and the weighted score can be used to select the top designs for *in vitro* expression and characterization.

**Figure 1.**
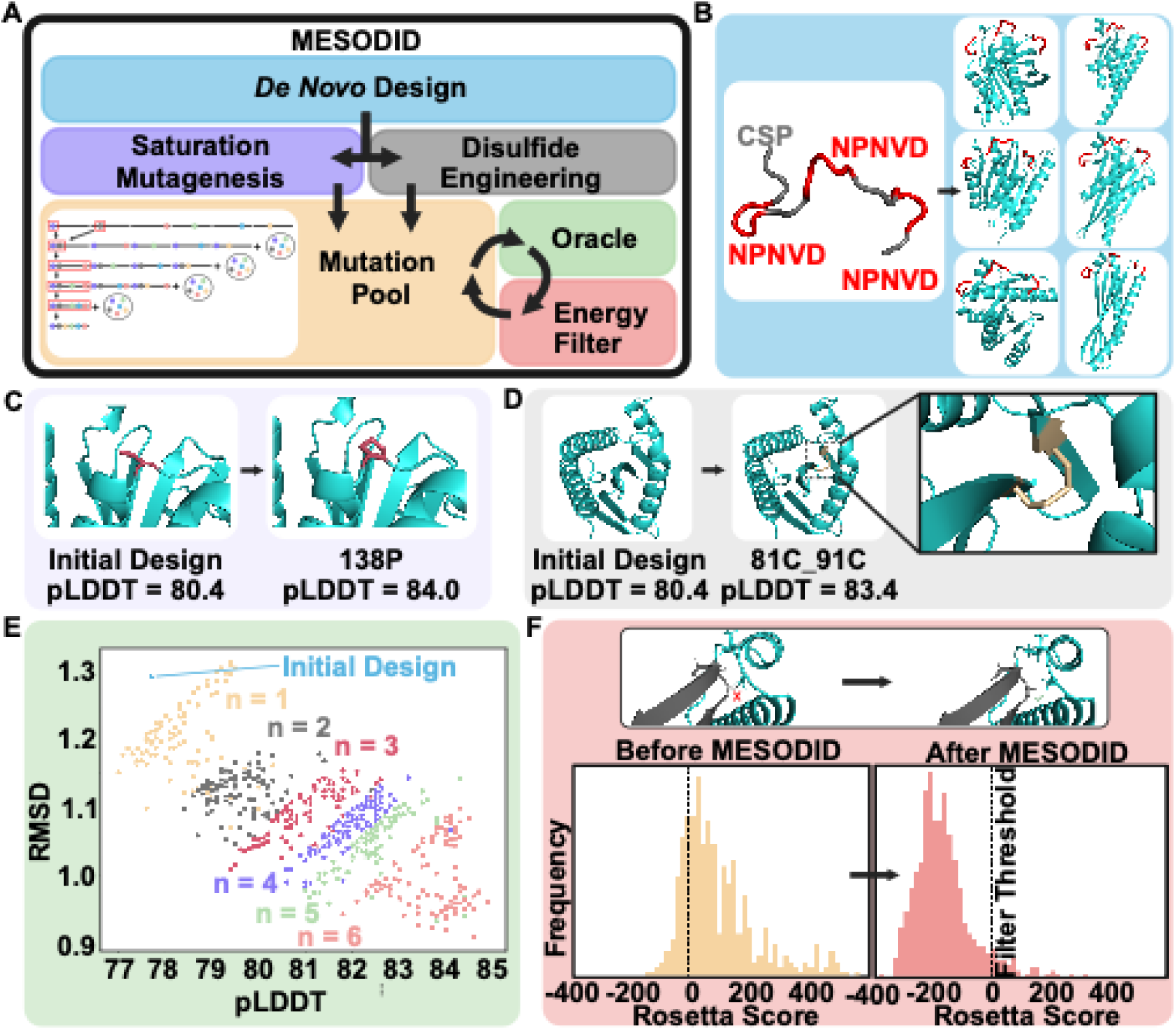
MESODID design pipeline. (A) MESODID combines deep learning protein design with multiple iterative refinement modules. Multiple refinement modules are used to generate a mutation pool, from which combinations of mutations are iteratively tested and filtered. (B) Depiction of structural motifs chosen for 3-NPNVD designs with examples of output scaffolds. (C) The first refinement module, saturation mutagenesis, mutates every amino acid outside the epitope of interest to every other possible amino acid. Inlaid graphic shows an example of a point mutation that improves pLDDT. (D) The second refinement module, disulfide engineering, identifies residue pairs that may be amenable to the introduction of a new disulfide bond based on patterns learned from a representative dataset from the PDB. Graphic shows an example of a new disulfide bond that improves pLDDT. (E) ESMFold is used as an oracle to predict the effects of newly introduced mutations. Graphic shows an example of improved computational metrics of top 100 mutation combinations tested for one backbone over subsequent rounds of refinement. (F) Designs pass through an energy filter before downselection to the next round of refinement. Inlaid graphic shows an example of improved Rosetta Score of immunogen-antibody complex over the course of MESODID.

### DESIGN, EXPRESSION, AND ANTIGENIC PROFILING OF TRIPLE MINOR REPEAT IMMUNOGENS

We employed MESODID to scaffold three copies of the minor repeat in one immunogen to accurately recapitulate the homotypic interactions observed between copies of L9 in complex with CSP (Figure 1B). In CSP, each NPNV minor repeat is followed by a DPNA tetrapeptide reported to exhibit limited direct interactions with L9 (2). However, upon further examination of the CSP-L9 paratope-epitope interaction, we noticed that aspartate in this DPNA tetrapeptide sits in a small pocket along with L9 CDRH3 aspartate D102. We further note that the aspartate of the tetrapeptide following the minor repeat has been reported having a large buried surface area similar to amino acids of the minor repeat upon complexation with L9 (3). We therefore included this aspartate in our epitope definition, where we focused on three discontinuous copies of NPNVD as each NPNVD motif comprises the full epitope bound by a single L9 Fab (2, 3)’. We initially generated 50,000 backbones each with 10 associated sequences for a total of 500,000 starting designs (Figure S3A). Those with a weighted score of less than 26.5 (equating to the top 19 designs), as predicted by ESMFold, subsequently underwent iterative rounds of refinement with approximately 1000 – 2000 new designs generated per backbone per round of refinement. Upon completing the iterative design steps of the pipeline, the top 20 refined designs for each backbone by weighted score as predicted by ESMFold were also predicted using Alpha Fold 2 (AF2) (35) (Figure S3B-D). Each of these designs had weighted score values of at most 23 (average pLDDT of 84.3, RMSD of 1.01 Å) and a Rosetta energy score of at most -350. Weighted scores correlated well between ESMFold and AF2 (Figure S3B-D). The design for each input backbone with the lowest weighted score after AF2 prediction was selected as the final design for validation in the lab, yielding 19 triple epitope designs for *in vitro* testing (Table S2). The designs were expressed using the same protocol as used for the single epitope designs. Remarkably, 16 of the 19 *de novo* designed proteins were successfully expressed to varying degrees of purity (Figure S4). Of the 16 designs, 9 showed SEC peaks at the expected elution volume for monomeric designs.

To assess whether these candidate immunogens were capable of binding L9, we performed ELISA, with the receptor binding domain of the middle east respiratory virus (MERS-RBD) as an unrelated negative control. All 16 expressed immunogens bound L9 to varying degrees, with no binding of L9 to the MERS-RBD observed (Figure S5). Critical to this design problem was the need to scaffold discontinuous epitopes such that L9 (or L9-like antibodies) would be able to bind the scaffold in such a way that the homotypic interactions between antibody copies would be maintained. To assess which designs were most likely to have successfully recapitulated these homotypic interactions, we performed an ELISA experiment to assess the binding of each design to a variant version of L9 with its homotypically interacting residues “knocked out” (L9_HOKO_). This variant was chosen based on previous data showing its ability to impair homotypic interactions between copies of L9, as well as its ability to impair L9’s protective efficacy in a malaria challenge model, and included F96Y_H_, D98A_H_, F28A_L_, R31A_L_, E68G_L_, and H70E_L_ (2). We hypothesized that our designs would be able to bind L9_HOKO_; however, designs with large changes in binding affinity between L9 and L9_HOKO_ would likely recapitulate L9’s homotypic interactions (Figure 2A). Two designs—design 4 and design 7—were especially promising, given their high change in binding affinity observed between L9_HOKO_ and L9.

**Figure 2.**
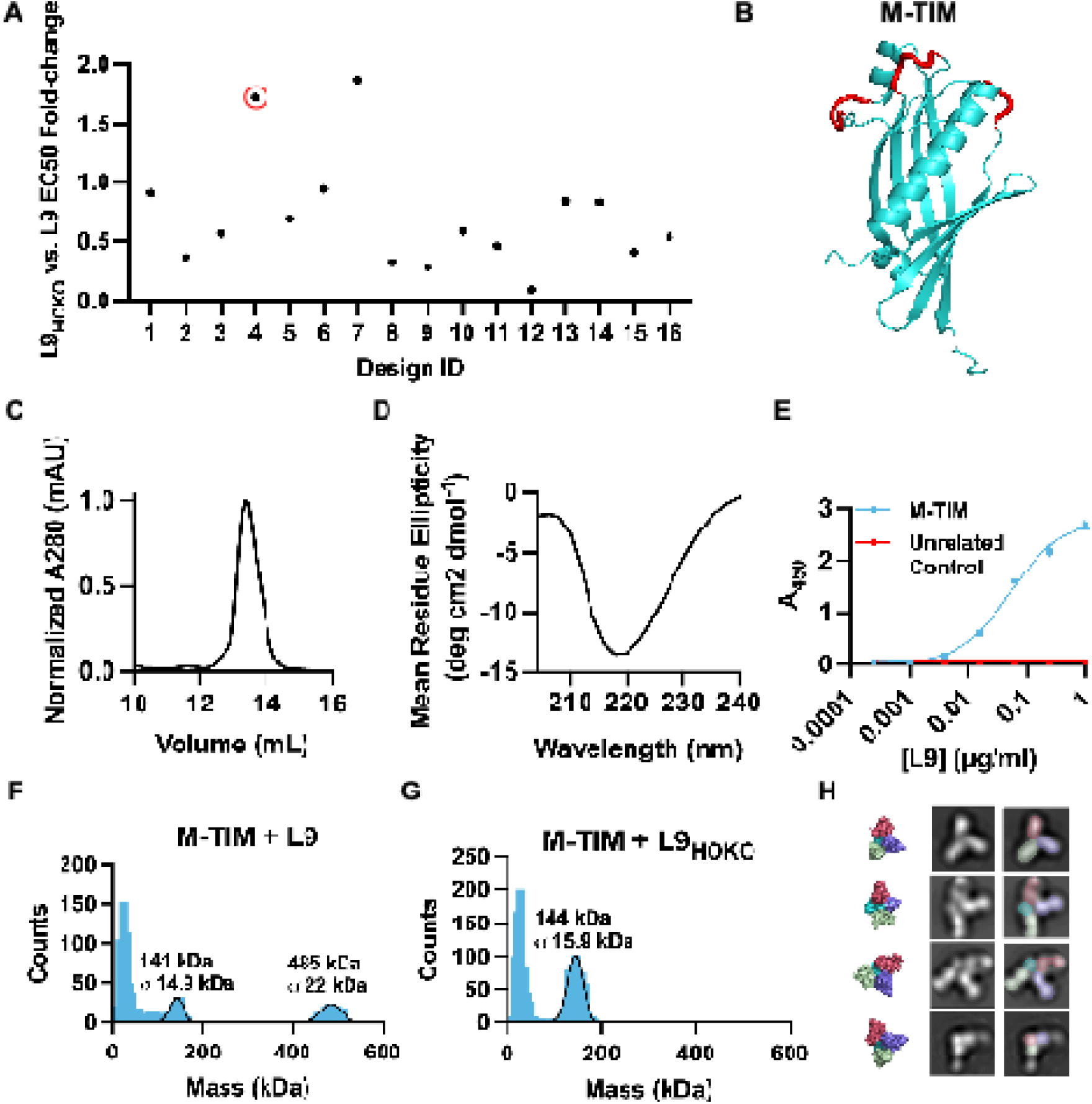
Biophysical characterization and antigenic profiling of top 3-NPNVD design M-TIM. (A) L9 vs L9_HOKO_ EC50 ratio of top 16 3-NPNVD designs. Red circle denotes top design downselected for further characterization (design 4). (B) AF2 prediction of M-TIM (design 4). Red residues represent the 3 NPNVD repeats. (C) SEC trace and (D) circular dichroism of M-TIM. (E) L9-M-TIM ELISA binding curve. MERS-RBD was used as an unrelated negative control. (F) Single particle mass photometry of M-TIM in complex with L9 and (G) L9_HOKO_. (H) Negative stain 2D classes of M-TIM in complex with L9. Individual fabs are shaded in green, dark pink, and purple, with M-TIM shaded in cyan. Model of three L9 fabs bound to M-TIM shown on the far left.

The AF2 structure prediction of design 4, which we will hereafter refer to as M-TIM, is shown in Figure 2B, with the three NPNVD motifs highlighted in red. M-TIM is 196 amino acids long, has an estimated molecular weight of 23 kDa and consists of a quasi-beta barrel morphology with an alpha helix running through the center. M-TIM expressed with a single, clean peak via SEC (Figure 2C), and circular dichroism confirmed a mostly beta secondary structure consistent with its AF2 prediction (Figure 2D). M-TIM’s ELISA binding curve showed high-affinity binding to L9 with an EC50 of 53 ng/mL (Figure 2E). Thus, M-TIM expressed well and had likely formed a well-folded protein core with high affinity to L9. However, while ELISA data suggested M-TIM may bind L9 in such a way that the homotypic interactions between copies of L9 was maintained, this data did not unequivocally demonstrate that M-TIM was capable of binding 3 copies of L9.

To determine the number of copies of L9 bound by M-TIM, we further evaluated M-TIM using single-particle mass photometry. Specifically, we complexed M-TIM with either L9 or L9_HOKO_ overnight and recorded mass distributions of the resulting complexes in solution the following day. Notably, M-TIM in complex with L9 showed a strong peak at the expected mass for a three-antibody-bound complex (expected: 473 kDa and observed: 485 ± 22 kDa) as well as a peak at the expected mass for either free antibody or one copy of antibody bound to M-TIM (expected IgG: 150 kDa; expected M-TIM: 23 kDa; observed: 141 ± 14.9 kDa), as these two species are indiscernible at the resolution of the assay (Figure 2F). In addition, complexing M-TIM with L9 Fab showed a mass distribution consistent with 3 copies of L9 bound to M-TIM (expected: 173 kDa, observed: 162 ± 25 kDa, Figure S6A-B). On the contrary, M-TIM in complex with L9_HOKO_ IgG showed a strong peak only at the expected mass for free antibody/one copy of antibody bound to M-TIM (observed: 144 ± 15.9 kDa, Figure 2G). This data suggested that M-TIM may orient 3 copies of L9 such that they are reliant on their homotypic interactions for their high binding affinity. To further confirm M-TIM’s ability to bind 3 copies of L9, we performed negative stain electron microscopy on the M-TIM-L9 Fab complex. Multiple 2D classes showed what appeared to be three copies of Fab bound to M-TIM (Figure 2H, Figure S6C). Taken together, this data suggests M-TIM is capable of binding three copies of L9 such that the homotypic interactions are important for the 3 to 1 stoichiometry. Design 7 was similarly characterized but appeared unable to successfully orient three copies of homotypically interacting L9 via mass photometry (Figure S7). Thus, M-TIM (design 4) was selected for further structural characterization.

### STRUCTURAL CHARACTERIZATION OF IMMUNOGEN M-TIM

To assess whether these homotypic interactions were recapitulated by M-TIM on a molecular level, we collected and analyzed single-particle cryo-electron microscopy (EM) data of the purified L9 Fab:M-TIM complex. In total, we collected 5,558 movies, picking ∼4.3 million particles with multiple 2D classes demonstrating three copies of L9 Fab bound in a trimeric fashion (Figure 3A, Figure S8). Using 434,079 high quality particles, we generated a 3D reconstruction of the complex at a resolution of 3.6Å. In the cryo-EM density map, we visualize three tightly packed L9 Fabs in complex with M-TIM (Figure 3B). The map features well-resolved density for all three L9 variable regions (Fv) as well as for M-TIM. We built the complex model using the previously published structure of CSP in complex with L9 (PDB: 8EK1) and the AF2 prediction of M-TIM as starting models (Figure 3C, Table S3). Of note, a predicted structure of the L9 Fab:M-TIM complex using AlphaFold 3 (36) resulted in each copy of L9 rotated and misaligned to the minor repeats, thus further validating the need for experimental structural characterization (Figure S9).

**Figure 3.**
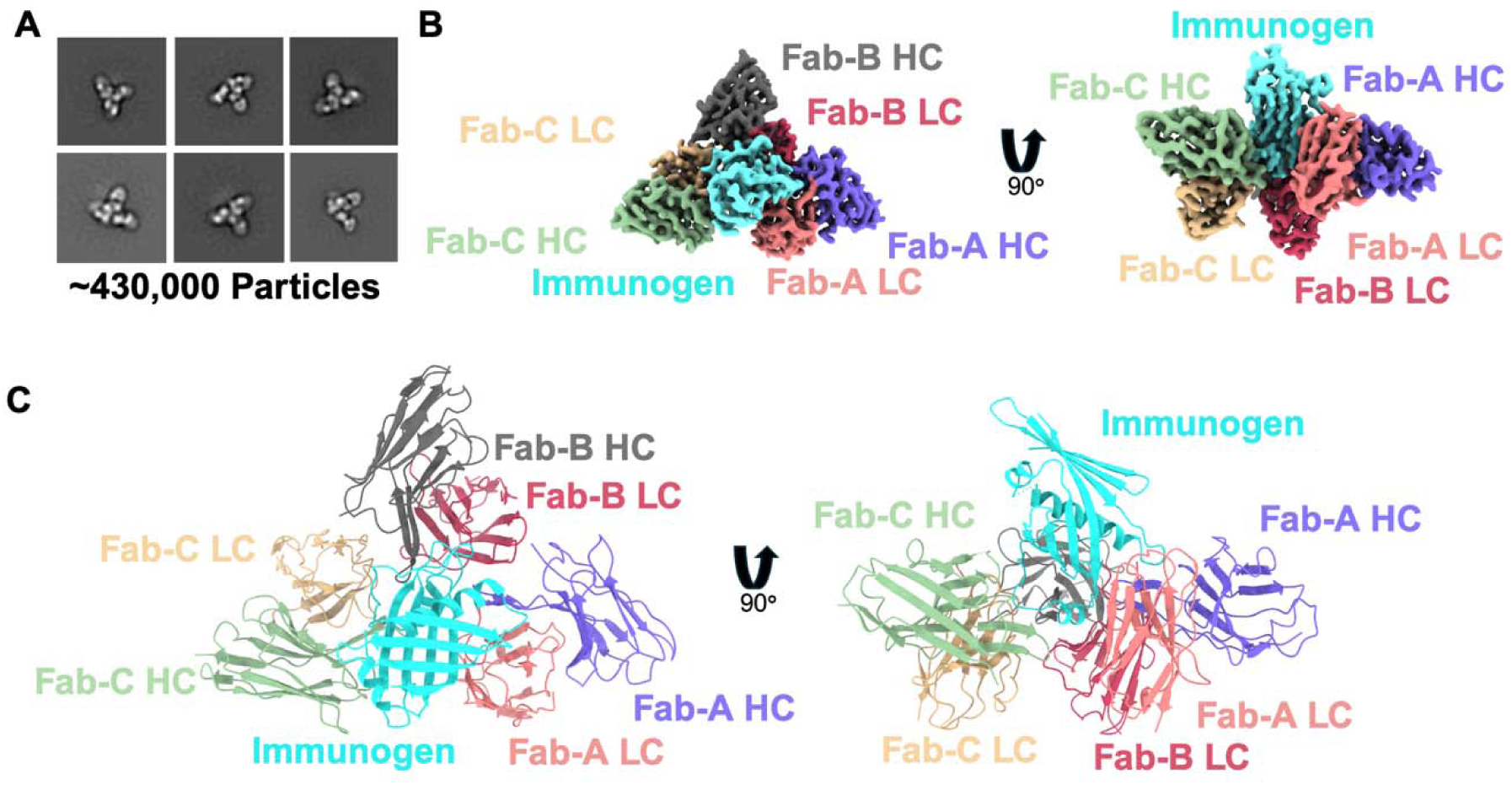
Cryo-EM structure of M-TIM in complex with L9. (A) Examples of 2D classes from cryo-EM dataset showing the trimeric M-TIM-L9 complex with total number of particles used for 3D reconstruction listed. (B) Electron density map of 3D reconstruction of L9-M-TIM complex. (C) Model of M-TIM-L9 complex.

The resulting structure demonstrates the desired topology of L9 CDRH3s engaging the NPNVD minor repeats. Importantly, L9 recognizes each of the three NPNVD motifs in M-TIM in an almost identical fashion to the way L9 recognizes wild type CSP’s minor repeats (Figure 4A, Figure S10). In addition, a detailed buried surface analysis confirms the importance of identical paratope residues in recognizing the minor repeats when compared to wildtype CSP (Figure 4B). In each epitope:paratope pair, N at the first position of NPNVD interacts with L9 CDRL1 W32_L_ hydrophobically and possibly by hydrogen bonding (Figure 4C). P at the second position of NPNVD packs hydrophobically with CDRL3 Y94_L_ (Figure 4D). N at the third position of NPNVD forms a hydrogen bond with CDRL3 R96_L_ (Figure 4E). V at the fourth position of NPNVD packs hydrophobically with CDRH3 Y97_H_ (Figure 4F). Finally, D at the fifth position of NPNVD forms a hydrogen bond with CDRH1 Y32_H_. Y32_H_ is positioned as a hydrogen donor for this hydrogen bond through its hydrophobic stacking with CDRH2 W52_H_ and CDRH1 Y32_H_ (Figure 4G) (2, 3)’.

**Figure 4.**
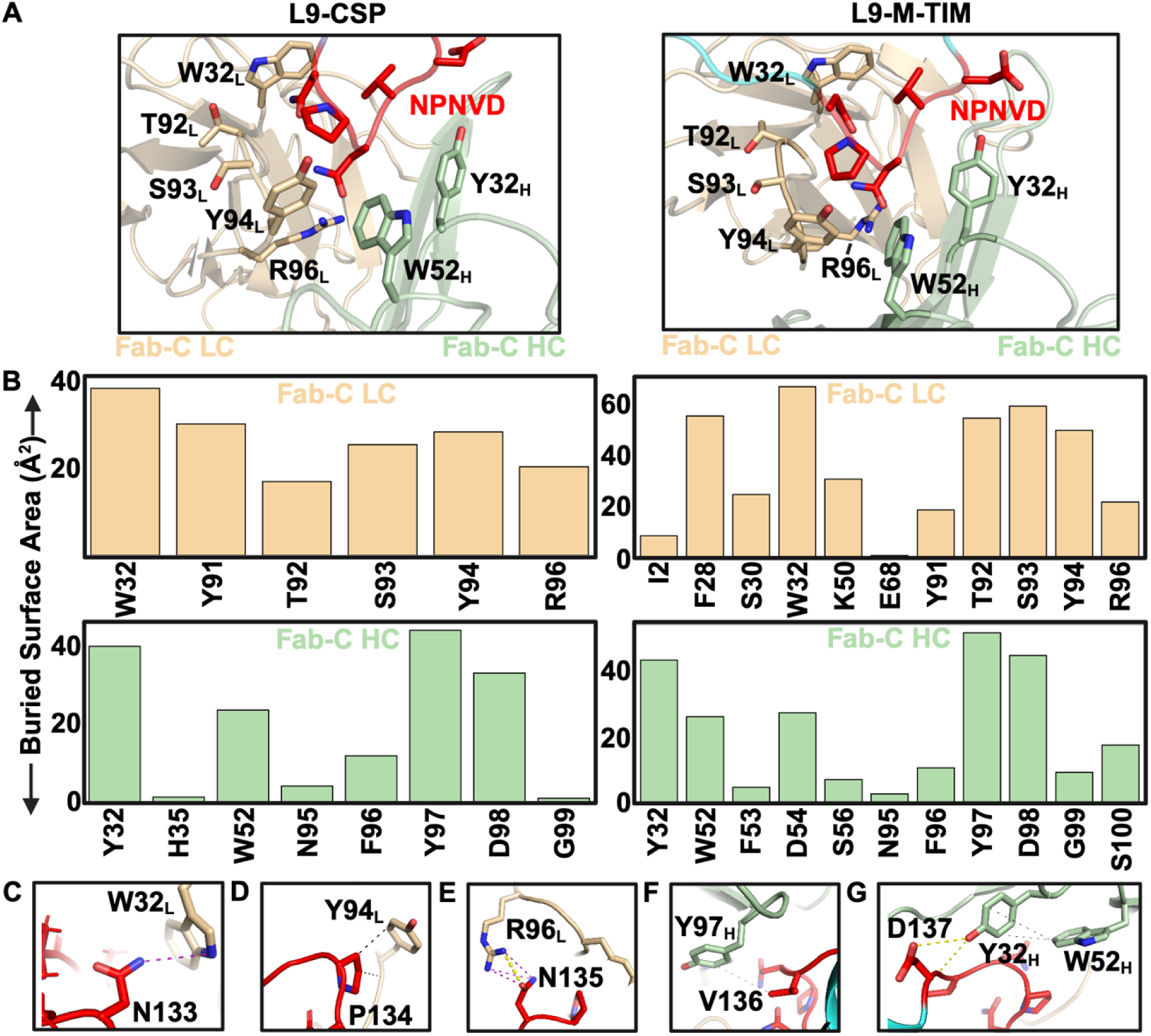
Structural comparison of epitope-paratope interactions between L9-CSP and L9-M-TIM complexes. (A) Representative interaction between NPNVD repeat scaffolded by M-TIM and L9 paratope. L9-CSP interaction is shown on the left with L9-M-TIM interaction on the right. (B) Buried surface analysis of important paratope residues involved in NPNVD recognition in L9-CSP vs L9-M-TIM complex. Example of (C) first asparagine of the NPNVD motif forming a hydrogen bond with W32_L_, (D) proline of the NPNVD motif interacting hydrophobically with Y94_L,_ (E) second asparagine of the NPNVD motif forming multiple hydrogen bonds with R96_L_, (F) valine of the NPNVD motif interacting hydrophobically with Y97_H_, and (G) aspartate of the NPNVD motif forming a hydrogen bond with Y32_H_, where Y32_H_ is positioned as a hydrogen donor through its hydrophobic stacking with W52_H_.

We further analyzed the area of L9 responsible for homotypic interactions. M-TIM orients the three copies of L9 such that the critical homotypic interactions between copies of L9 are maintained (Figure 5A,B) with a total homotypic-interface C_α_-RMSD of 1.2 Å (Figure S10). The Fab-A:Fab-B and Fab-B:Fab-C homotypic interfaces are nearly identical. In the cryo-EM structure, extensive hydrophobic and polar contacts contribute to the stability of these interfaces, specifically between the CDRL3 and light chain FR3 of Fab-C with the CDRH1, CDRH3, and heavy chain FR1 of Fab-B. F27_H_, F28_L_, F96_H_, and F100C_H_ form key interacting residues as a part of a stabilizing hydrophobic pocket along the interface, as noted in the recently published CSP-L9 cryo-EM structures (2, 3)’. R31_L_ from the CDRL1 of Fab-C additionally forms key interactions with F100C_H_ within the CDRH3 of Fab-B. H70_L_, a framework residue, interacts extensively with Q1_H_ of the Fab-B CDRH1, and E68_L_ forms extended contacts along the homotypic interface. Together, these residues have been previously shown to be required for the stability of the homotypic interface, as well as the protective efficacy of L9 *in vivo* (2, 3).

**Figure 5.**
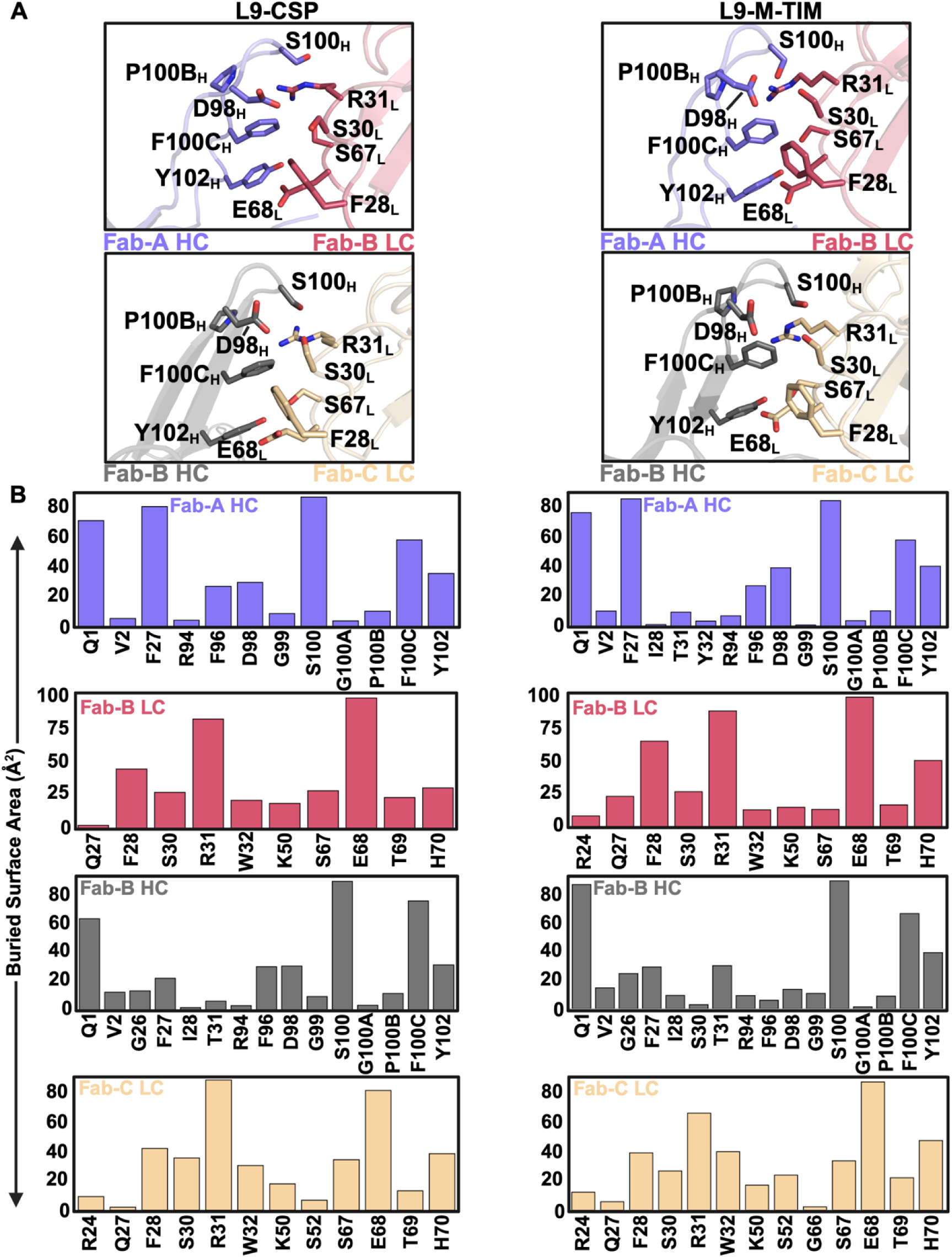
Comparison of homotypic interactions between copies of L9 Fab in L9-CSP and L9-M- TIM structures. (A) Interactions between homotypically-interacting residues in L9-CSP vs L9-M- TIM complex. (B) Buried surface analysis of important homotypically-interacting residues in L9- CSP vs L9-M-TIM structures.

One of the key hypotheses motivating this work is that scaffolding the minor repeats will structurally stabilize the repeats such that they occupy states capable of binding L9-like precursors for a higher percentage of time than they would if administered as free peptide. To test this hypothesis *in silico*, we performed 3 x 1 µs molecular dynamics simulations at 37°C of either M-TIM or the 27 amino acid stretch of wild type CSP that contains 3 copies of the minor repeat, labelled as 3 NPNVD Peptide (Figure 6). In all simulations, the 3 NPNVD Peptide or M-TIM began with the three NPNVD repeats in the same configuration in which L9 recognizes them. Under these conditions, the free peptide was highly dynamic, with the RMSD of the minor repeats to the reference structure quickly jumping to between 6 and 12 Å (Figure 6A,B). The RMSD of the minor repeats remained highly volatile across the duration of the simulation time, never coming close again to adopting a conformation similar to its starting pose. M-TIM, however, demonstrated impressive stability of the minor repeats across three independent simulations, with the total RMSD across all three repeats consistently maintained near 2 Å from the starting pose (Figure 6C,D). These simulation results support the hypothesis that M-TIM can more stably present 3 NPNVD motifs than a free CSP peptide.

**Figure 6.**
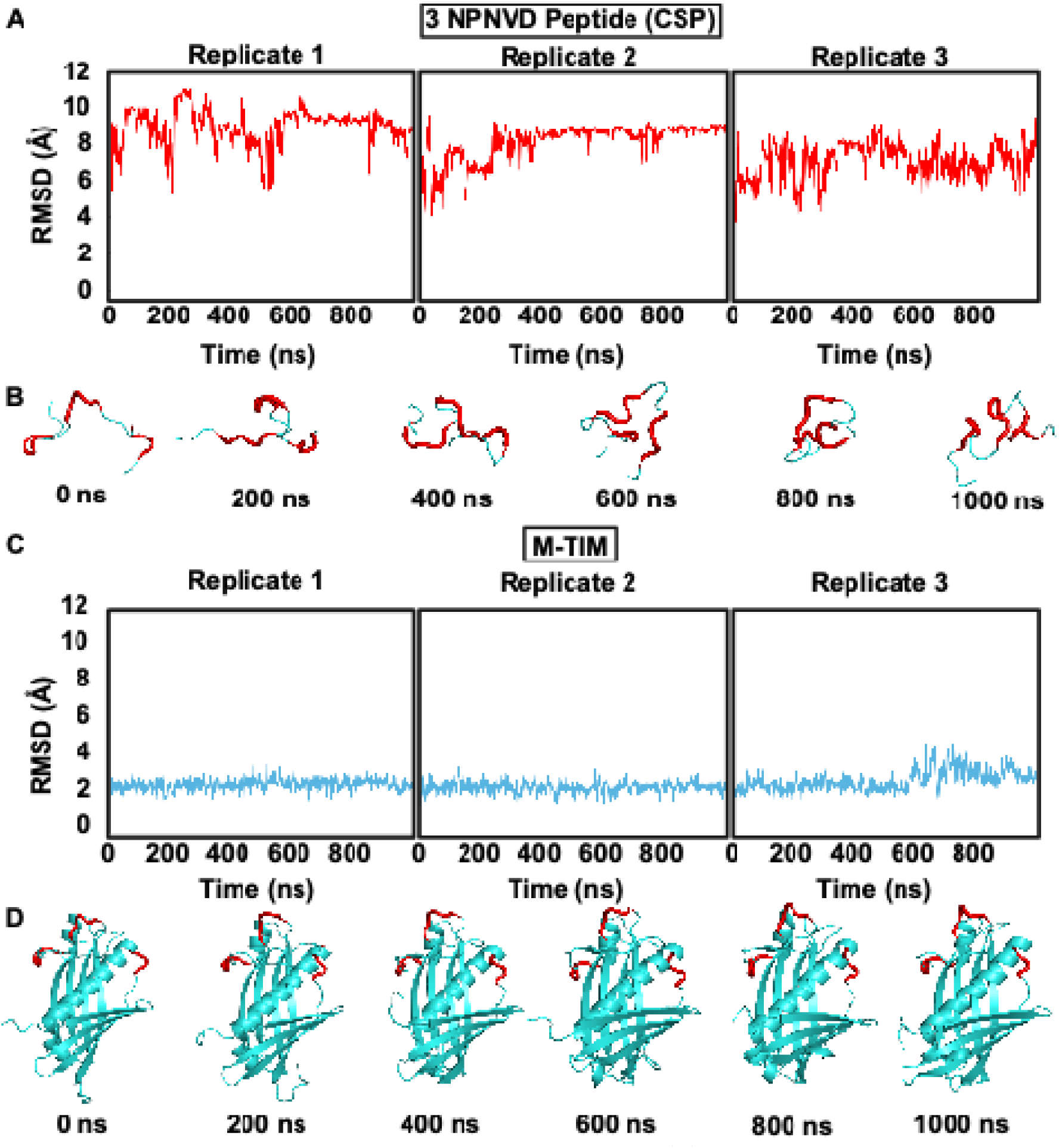
Molecular dynamics simulations of CSP vs M-TIM. (A) RMSD is shown for 3 NPNVD repeats in CSP-based peptide for 3 independent 1 µs simulations. (B) Structural snapshots of CSP-based peptide are shown from one simulation. (C) RMSD is shown for 3 NPNVD repeats in M-TIM for 3 independent 1 µs simulations. (D) Structural snapshots of M-TIM are shown from one simulation.

## Discussion

In this study, we successfully designed immunogens that present both one and three copies of *Pf*CSP’s minor repeat. Our top three-repeat-based design, M-TIM, stabilizes 3 discontinuous copies of the *Pf*CSP minor repeat, such that each copy of the minor repeat binds L9 while maintaining homotypic interactions between the antibodies critical for L9’s protective efficacy. To accomplish this, we developed MESODID, an iterative deep-learning based, structure-guided protein design pipeline. MESODID could similarly be applied to other repetitive epitope problems, of which many are known within bacterial and viral pathogens (37–41). In particular, the HIV-1 envelope, Sars-CoV-2 spike, and influenza hemagglutinin (HA) proteins contain multiple repeated neutralizing epitopes across their trimeric structures (39–41). For example, the monoclonal antibody CR9114 targets a conserved epitope in each copy of the stem of the HA trimer and has been shown to protect broadly across multiple influenza A and B viruses (42). Stabilizing these epitopes as a part of an immunogen could help drive a more broadly neutralizing antibody response as opposed to current influenza vaccination strategies. Of note, using the MESODID pipeline led to a high experimental success rate and required no additional *in vitro* optimization via yeast or mammalian display. A cryo-EM structure of M-TIM in complex with L9 confirms M-TIM’s successful recapitulation of homotypic interactions between copies of L9. Importantly, molecular dynamics simulations corroborate the marked structural stability of the minor repeats scaffolded by M-TIM, in comparison with a free peptide containing three copies of the minor repeat. These data suggest minor repeats within M-TIM may occupy states capable of binding L9-like antibodies for a higher percentage of time than they do when presented via free peptide. Together, these results unveil a designed immunogen that successfully replicates the repetitive structure of key epitopes and drives crucial homotypic interactions of the target antibodies.

Currently approved vaccines for malaria elicit variably potent antibodies, which is one of the aspects thought to contribute to their imperfect protection (1). These vaccines present 19 copies of the major repeat, and, of note, do not incorporate the minor repeat. In solution, these repeats appear to be highly dynamic, rarely adopting conformations capable of binding homotypically interacting antibodies such as L9 (43). Given its structural stabilization of the minor repeats, M-TIM could be used in a next-generation CSP-targeted vaccine with an improved ability to induce homotypically interacting L9-like antibodies, thus providing improved protection over state-of-the-art CSP-based subunit vaccines. Additionally, M-TIM could be used in tandem with RTS,S or R21 to generate additional synergistic immunity with concomitant improvement in protection. Given the high structural similarity between the stabilized secondary structure induced by highly potent anti-NANP/NPNA antibodies and the structural motif recognized by L9, M-TIM is expected to “shape” the anti-NANP/NPNA immune response toward more potent, homotypically-interacting antibodies (26).

Another potential application for M-TIM is the discovery of improved antibodies against the minor repeats of CSP. Recent landmark clinical trials have demonstrated marked protective efficacy in preventing clinical malaria by using monoclonal antibodies as a form of passive immunization (44, 45)’. These antibodies were first isolated from people who were vaccinated with radiation-attenuated sporozoites by using CSP, major repeat, or junctional region-based probes (17, 18)’. In similar fashion, M-TIM could be used as a probe to discover more potent, homotypically interacting antibodies from the sera of people previously vaccinated with radiation-attenuated sporozoites. Novel, potent, anti-CSP antibodies could also be discovered by vaccinating with M-TIM in a humanized mouse model in which the vast majority of B cells express the inferred germline precursor of L9. This has previously been done for CIS43, another highly potent anti-CSP antibody, resulting in the discovery of antibodies significantly more potent than CIS43 (46). In conclusion, we have developed MESODID, a powerful tool to design immunogens targeting repeating epitopes, and have used it to engineer M-TIM, a promising malaria vaccine candidate that uniquely recapitulates critical homotypic antibody interactions – a strategy with broad implications for vaccine design against diverse infectious diseases.

## Materials and Methods

### Computational Design

MESODID begins with selection of a structural motif for scaffolding. In this case, the three NPNVD motifs present in chain P of PDB 8EK1 were selected. RFdiffusion (27) was then used to generate 50,000 protein backbones capable of presenting the 3 selected NPNVD motifs, with 30-60 amino acids on both the N- and C-terminus of the designs as well as between motifs. Preliminary runs of 100-1000 designs across varied topologies connecting NPNVD motifs were run to assess the proper number of amino acids between motifs in order to form a well-packed protein without generating an unnecessarily large immunogen. Only the topology with motifs connected in sequential order was chosen for the larger run of 50,000 backbones, as this led to designs with the closest epitope similarity to PDB 8EK1 as measured by C_α_ root mean square deviation (C_α_-RMSD). ProteinMPNN (28) was then used with a sampling temperature of 0.1 to generate 10 sequences per RFdiffusion backbone for a total of 500,000 designs. Designs were subsequently subjected to structure prediction using ESMfold (29) and filtered based on a weighted score, given by *weighted score = 10 * RMSD + 0.6 * (100 – pLDDT)* where pLDDT is the protein local distance difference test (pLDDT) and RMSD is the root mean square deviation of the scaffolded epitope to the structural epitope described from PDB 8EK1 (2).

The top designs underwent iterative rounds of refinement to improve the designs’ weighted score. To begin the refinement process, each design underwent two refinement modules: saturation mutagenesis and disulfide engineering. In the saturation mutagenesis module, all possible single mutations outside the epitope of interest were structurally predicted using ESMFold. Designs that improved the weighted score were down-selected for further rounds of refinement. In the second module, a previously published disulfide scanning code (30) was used to search for 3D geometries and amino acid environments capable of introducing stabilizing disulfide bonds to each top design. Those designs with newly introduced disulfide bonds that improved the weighted score were similarly down-selected for further rounds of refinement. Mutations from both modules that improved these computation metrics were put into an initial “mutation pool.” From this mutation pool, combinations of mutations were tested.

To determine whether combinations of mutations would be selected for subsequent rounds of refinement, we used two main optimization metrics. The first was the weighted score. This score combined both pLDDT and RMSD into one score, while emphasizing the greater importance of RMSD to our particular design problem.

The second metric was determined first by aligning the predicted structure to the structure of 3 homotypically interacting copies of L9 from PDB 8EK1. This complex was then subjected to fixed-backbone relaxation using the Relax protocol in Rosetta (31–34)’. The Rosetta Score post-relaxation, which would reflect any energetically unfavorable interactions that may prevent the ability of 3 copies of L9 to bind, was used as a second metric for further optimization.

Designs that passed both weighted score and Rosetta score filters (a new weighted score lower than the top weighted score from the previous round of refinement and a Rosetta Score post-relax of less than zero) were placed into a “passed” category. The top 100 combinations of mutations from each round were selected from this passed category for further optimization. These combinations of mutations were then themselves combined iteratively in the next round of refinement. This proceeded for 4-6 rounds of refinement/design.

Thus, at the end of refinement, designs had been optimized both for their computational probability of folding accurately (pLDDT) and presenting the correct epitope (RMSD), but also for the energetic favorability of simultaneously binding three copies of antibody without introducing energetically unfavorable clashes between the antibodies and the candidate immunogen (Rosetta score post-relaxation). Top designs from ESMFold were finally validated using AF2. Optimized designs that performed similarly between ESMFold and AF2 were chosen as final constructs to be ordered and tested in the lab.

### Protein expression and purification

DNA constructs containing the output sequences from RFdiffusion and ProteinMPNN were cloned into the pVax vector with N-terminal Kozak and IgE leader sequences as well as a C-terminal 6xHisTag. Constructs were codon optimized for expression in *homo sapiens* by an optimization tool (Genscript). These constructs were transiently transfected into Expi293F cells using the cationic lipid-based ExpiFectamine™ 293 Reagent with supernatant harvested 6-7 days later. The supernatants were subsequently purified via affinity chromatography using an IMAC nickel column on an AKTA Pure 25 purification system (Cytiva). Appropriate fractions from affinity chromatography were pooled, concentrated, and dialyzed with 1X PBS before further purification using size exclusion chromatography on a Superdex 75 Increase 10/300 column. Fractions were pooled and concentrated to ∼1 mg/mL before being flash frozen in liquid nitrogen and stored at - 80C pending further experimental characterization. Antibodies were similarly cloned into pVax, with two separate constructs for the respective heavy and light chains. Both constructs were co-transfected in equal parts to Expi293F cells and purified using a Protein A column and the AKTA Pure 25 purification system. Appropriate fractions were pooled and stored at 4C until use.

### ELISA

To test binding of designed antigens to L9 antibody, high binding 96-well Flat-Bottom, Half-Area Microplates (Corning) were coated with 90 nM (1.9 - 2.3 µg/mL) antigen at 4C overnight. The next day, plates were blocked with 5% milk/1x PBS/0.01% Tween-20 at room temperature for ≥ 1 hour at ambient temperature. Plates were incubated with serially diluted primary antibody for 1 hour, followed by HRP-conjugated goat anti-human IgG Fc antibody for 1 hour, all at ambient temperature. Finally, plates were incubated with 1-Step^TM^ Ultra TMB-ELISA Substrate Solution for 5 minutes, after which the reaction was quenched with 1 M H_2_SO_4_. In between each step, plates were washed with 1x PBS/0.05% Tween-20. Absorbance of samples was read at both 450 and 570 nm, with the latter being subtracted from the first for each well. Background absorbance was also measured from blank wells and subtracted from each well before analysis, which was performed in Prism 10.

### Circular dichroism

Samples of SEC-purified protein were concentrated to ≥ 1 µM in Na_3_(PO_4_) to a volume of ≥ 300 µL and loaded into a 1 cm path-length cuvette. Cuvettes had previously been washed with deionized water, followed by 70% ethanol and allowed to dry. Cuvettes were then loaded onto either an Aviv 202 Circular Dichroism Instrument or a Jasco Model J-815 Circular Dichroism Spectrometer. CD spectra were collected in either 1 or 2 nM intervals starting at 240 nm for 60 seconds at each step.

### Mass photometry

Samples were first diluted in fresh, 0.2 µm-filtered 1X PBS to 200 nM. For complexing assays, antibody and were mixed in a 1:1 molar ratio and allowed to come to equilibrium at 4°C overnight. Samples were then brought to a final concentration of ∼20 nM by diluting ∼1:10 in fresh PBS directly on the stage of a Refeyn TwoMP mass photometer. Samples were loaded onto Refeyn sample well casettes which were adhered to Refeyn MassGlass UC slides. Counts were acquired for 60 seconds using the Acquire^MP^ software. A gaussian distribution was fit to the count distribution using Discover^MP^ software and plotted using Prism 10.

### M-TIM-L9 complex purification

To form the L9 Fab:M-TIM complex, a > 7-fold molar excess of L9 Fab was incubated with M-TIM at 4 C overnight in TBS. The following day, ∼500 µL of complex was purified via SEC on a Superdex 200 Increase 10/300 GL column. Appropriate fractions (Supplemental Figure 6) were concentrated and flash frozen before storage at -80 C.

### Negative stain electron microscopy collection and processing

Purified L9 Fab:M-TIM complex was diluted in TBS to ∼0.02 mg/mL. A total of 3 µL sample was adsorbed onto glow-discharged carbon-coated Cu400 EM grids. Grids were stained twice with 3 µL of 2% uranyl formate. The first stain was immediately blotted off onto filter paper after administration of stain, followed by addition of a second round of stain for 90 seconds. This last stain was then blotted onto filter paper. Images were collected on a FEI Tecnai T12 microscope with a Oneview Gatan camera at 67200x magnification and a pixel size of 2.356 Å. Data processing was performed with Relion v5.0.

### Cryo-EM sample preparation

Purified L9 Fab:M-TIM complex was diluted to ∼0.05 mg/mL. UltrAuFoil 1.2/1.3 grids (Quantifoil) were glow discharged and then used immediately for graphene oxide application prepared using a previously published protocol (47). Graphene oxide-coated grids were blotted for 4 seconds with 4 uL of sample at 4C and 100% humidity before being plunge frozen into liquid ethane using a Vitrobot Mark IV. Grids were stored in liquid nitrogen until imaging.

### Cryo-EM data collection

Automated data acquisition was performed using aberration-free image shift (AFIS) protocol through the EPU software (ThermoFisher). Dose-fractionated micrographs were collected on a Thermo Fisher Glacios microscope equipped with a Falcon 4 detector using a total dose of 50 e-/ Å^2^ across 49 frames and a defocus range of -0.8 to -2.2 µm, at 150,000x magnification and with 0.9475 Å pixel size. A total of 5,558 movies were collected for the final reconstruction.

### Single particle cryo-EM data processing

Data processing was performed in Relion (v5.0.0) and CryoSPARC (v4.3.1) according to standard cryo-EM data processing workflows. Briefly, motion correction was performed in Relion using MotionCor2, and the contrast transfer function (CTF) was estimated with the Patch CTF estimation tool in CryoSPARC. The CryoSPARC blob picker was then used to pick circular particles, resulting in a total of ∼4.3 million particles picked. These particles underwent multiple rounds of 2D classification and select 2D classes were manually picked (comprising a particle stack with 116,287 particles) to undergo ab-initio 3D reconstruction, resulting in a 3D class corresponding to M-TIM bound to 3 copies of L9. This class, along with several 3D classes reconstructed from noisy 2D classes, was used to perform several rounds of hetero refinement on a larger group of 1,546,252 selected particles, resulting in 434,079 particles in the highest resolution 3D class. These particles underwent several rounds of Bayesian polishing in Relion, followed by non-uniform and local refinement in CryoSPARC, resulting in the final 3.6 Å density map. The auto-sharpened map from cryoSPARC was further processed using EMReady (48) with default parameters.

### Atomic Model Building

The cryo-EM structure of L9 in complex with recombinant shortened CSP (rsCSP; PDB 8EK1) was used along with the AF2 prediction of the M-TIM structure to generate an initial model. The model was docked into the M-TIM:L9 density map in Phenix and refined using Coot and Phenix refinement. Model-to-map fit was validated using Phenix and the wwPDB validation server.

### Structural analysis

Buried surface area calculations were performed using UCSF ChimeraX. Figures depicting structural features were generated in UCSF ChimeraX or PyMol.

### Molecular dynamics simulations

All-atom MD simulations were performed using GROMACS v2023.2 (49) using the AMBER99SB- ILDN protein force field (50) with the TIP3P (51) water model. Coordinates were output every 10 picoseconds with an integration time step of 2fs. A Verlet cutoff scheme (52) was used for short-range interactions while Particle Mesh Ewald (PME) (53) was used for long-range interactions. Simulations were performed in cubic simulated boxes with 10 angstrom padding, and periodic boundary conditions were used in all three cartesian dimensions. Temperature and pressure were held constant at 310 K and 1 bar, respectively (NPT). Temperature was held constant with a velocity-rescaling thermostat, while pressure was held constant by employing an isotropic Berendsen barostat (54), with a time constant of 0.5 picoseconds and an isothermal compressibility of 4.5 x 10^-5^ bar^-1^. The primary input was either free 27mer CSP-based peptide from PDB 8EK1 or the AF2 prediction of M-TIM. Structures were prepared using Free Maestro and properly solvated to neutral charge with sodium and chloride ions before initiation of simulation runs. Three independent trajectories for both 3 NPNVD CSP peptide and M-TIM were acquired for 1 μs each using the Wistar High Performance Computing Cluster. For analysis, 1000 equally spaced frames were extracted from each trajectory using gmx trjconv, and gmx rms was used to compute RMSD of the 3 NPNVD repeats to their initial position in PDB 8EK1.

## Acknowledgments

We would like to thank Ashok Nayak and the Cryo-Electron Microscopy Facility at Thomas Jefferson University for providing training and access to the Thermo Fisher Glacios Electron Microscope. JP was supported by Wistar Institute startup funds. AOJ was supported by NIH/NIAID R01AI103280 and the Doris Duke Charitable Foundation Paragon of Research Award.

## Data, Materials, and Software Availability

Cryo-EM map was deposited in the Electron Microscopy Data Bank under accession number EMD-49059. Atomic coordinates have been deposited into the PDB under accession number 9N6E. All other data are included in the manuscript and/or supporting information.

