## Supplementary Material for "Machine learning enables *de novo* multi-epitope design of plasmodium falciparum circumsporozoite protein to target trimeric L9 antibody"

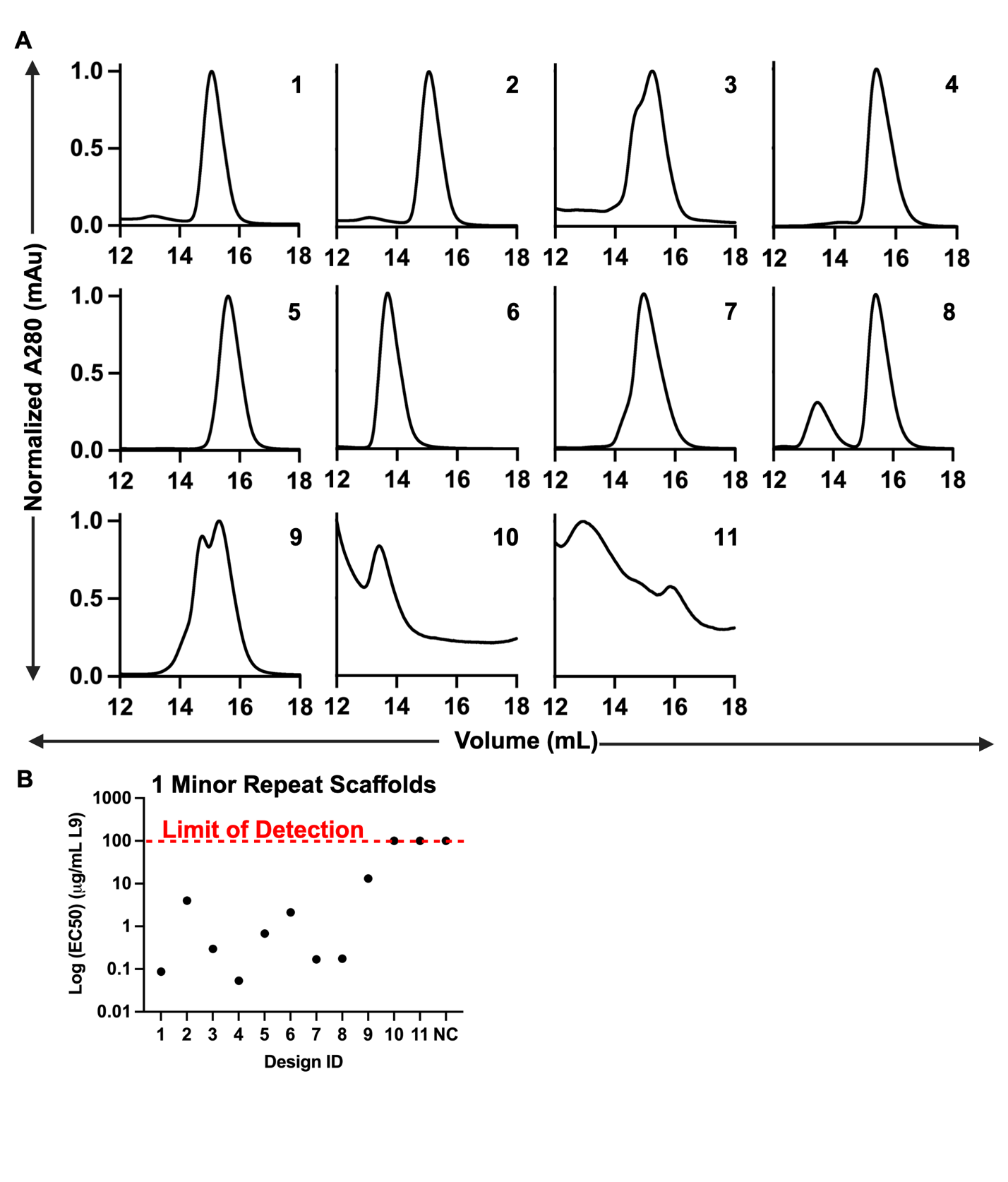

**Figure S1.**  Expression and antigenic profiling of top 1-NPNVD designs. (A) SEC trace for top 11 1-NPNVD design. (B) EC50 values for top 11 1-NPNVD designs. Red dashed line depicts limit of detection at 100 µg/mL. NC depicts the negative control, an HIV-1 Envelope trimer.

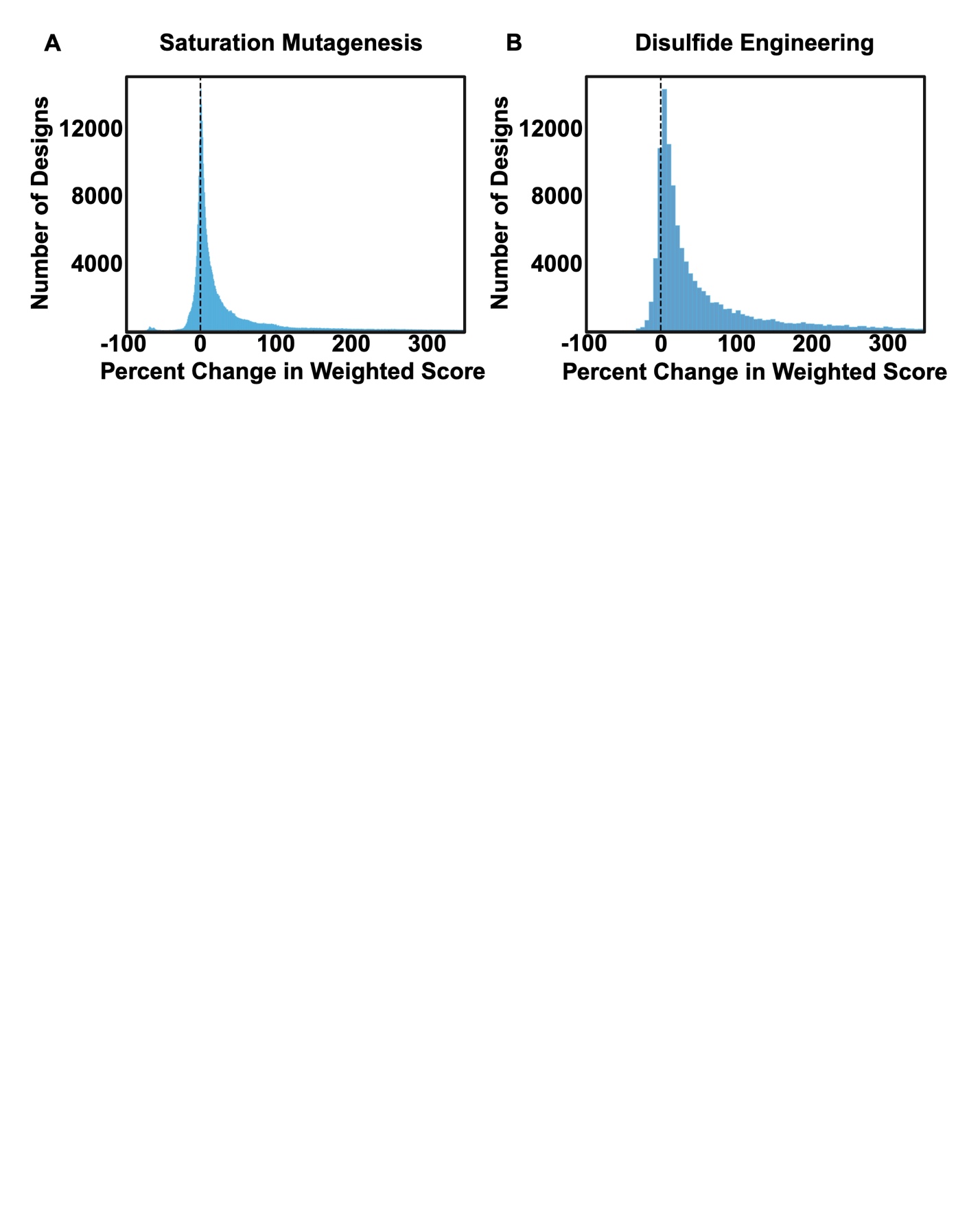

**Figuree S2.** Improvement of single mutants on weighted score. (A) Single mutations from saturation mutagenesis module and (B) disulfide engineering modules are plotted as the percent change in weighted score from the original design. 24.1 % of saturation mutagenesis mutations improved the weighted score from the original design while 11.6% of new disulfides improved weighted score.

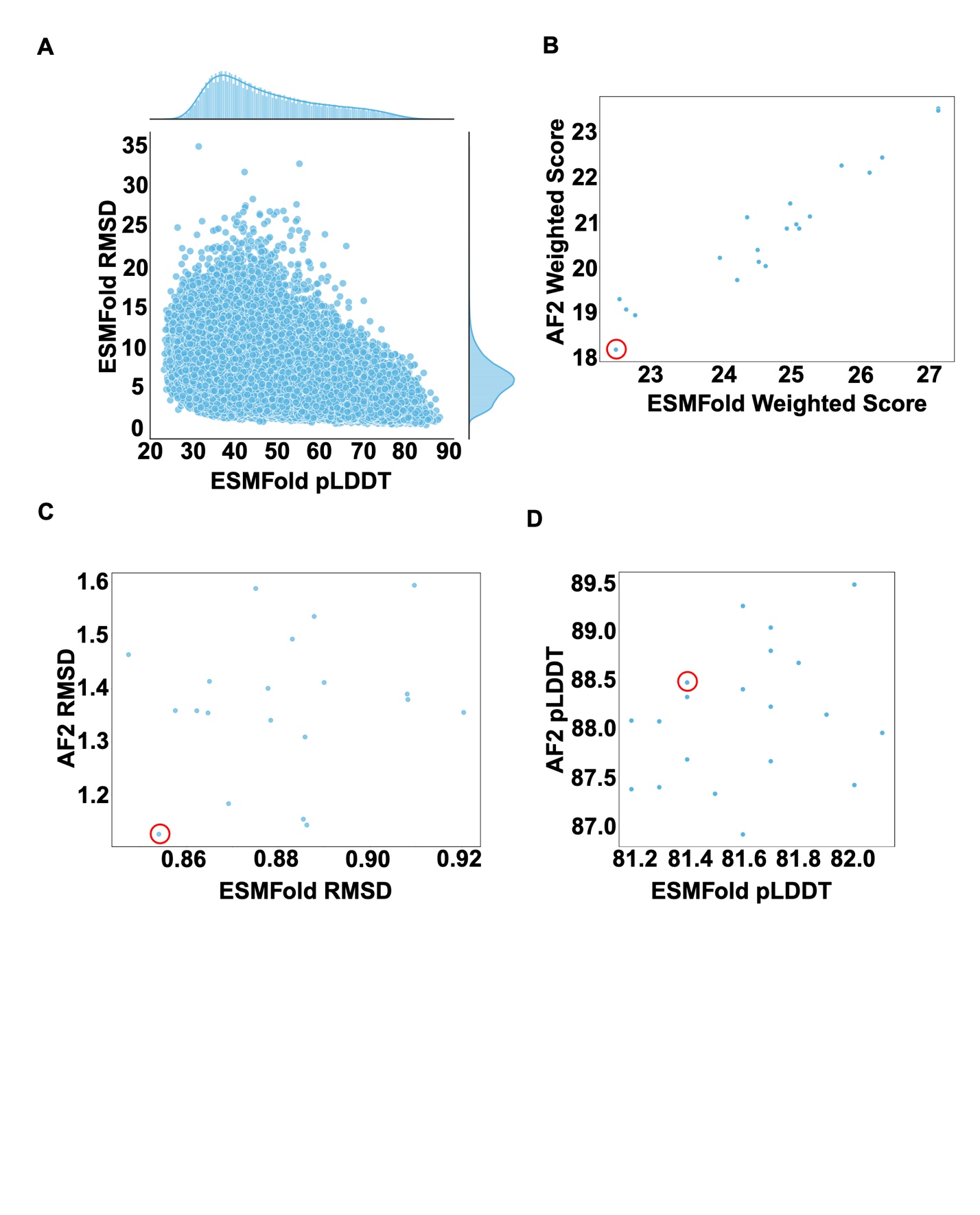

**Figure S3.** Computational design of M-TIM. (A) Initial output from de novo design with ESMFold as an oracle. (B) ESMFold weighted score vs AF2 weighted score of top 20 permutations of M-TIM design. (C) ESMFold RMSD vs AF2 RMSD of top 20 permutations of M-TIM design. (D) ESMFold pLDDT vs AF2 pLDDT of top 20 permutations of M-TIM design.

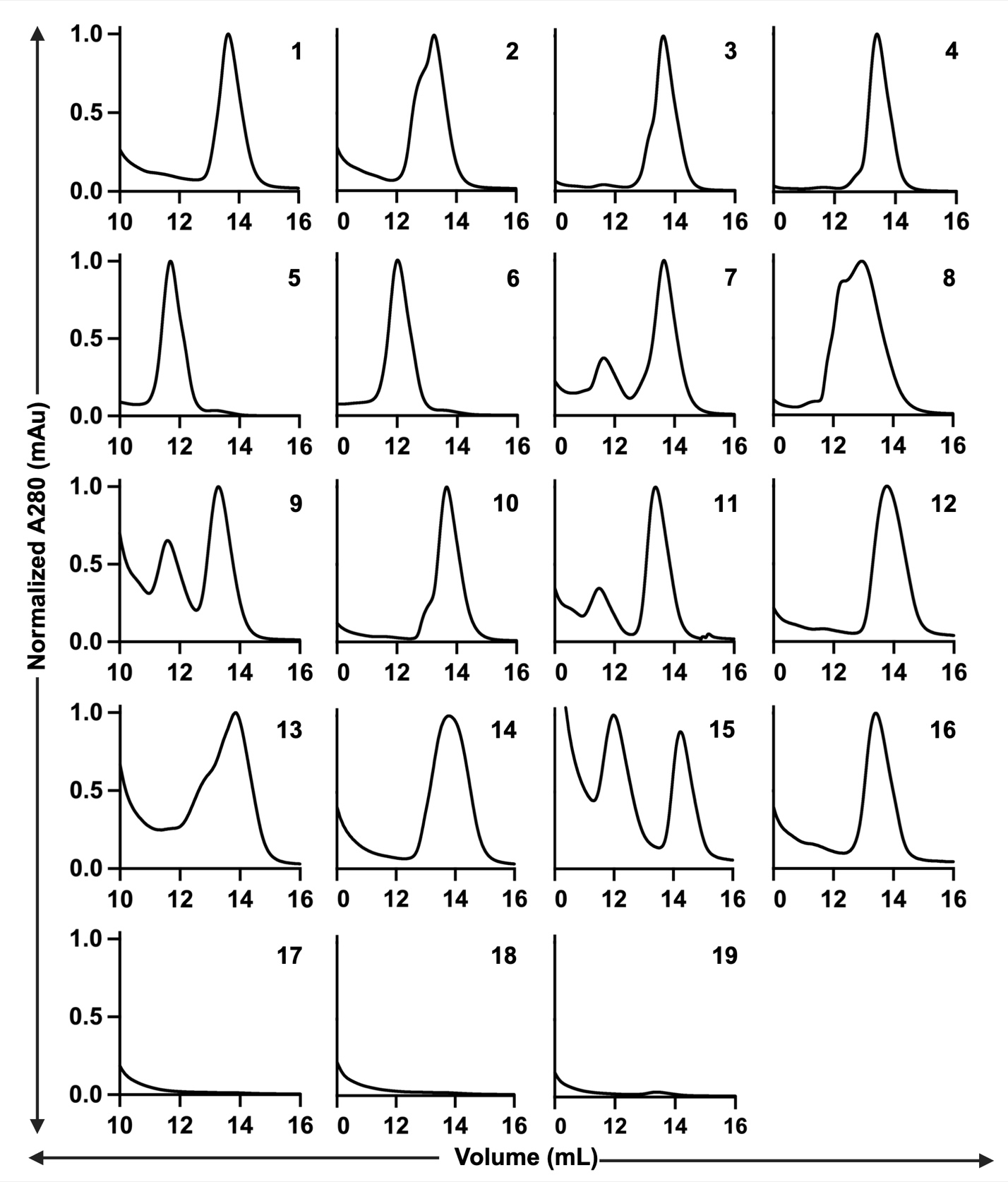

**Figure S4.** SEC traces of all 19 transfected 3-NPNVD designs.

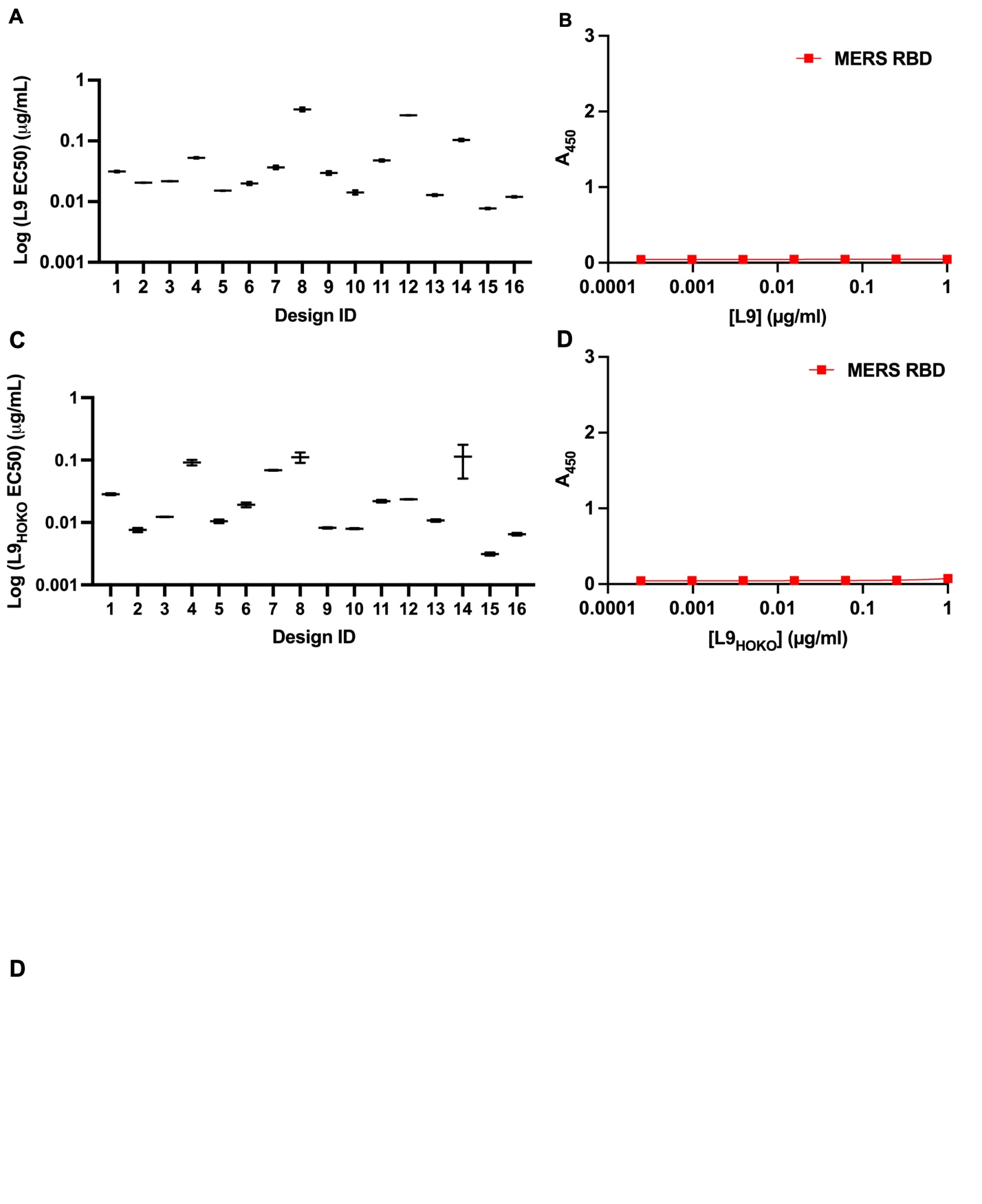

**Figure S5**. Antigenic profiling of top 16 3-NPNVD designs. (A) EC50 values for top 16 3-NPNVD designs binding to L9. (B) ELISA binding curve for MERS-RBD binding to L9. (C) EC50 values for top 16 3-NPNVD designs binding to L9_HOKO_. (D) ELISA binding curve for MERS-RBD binding to L9_HOKO_. For all experiments, error bars represent SEM for n=2 replicates.

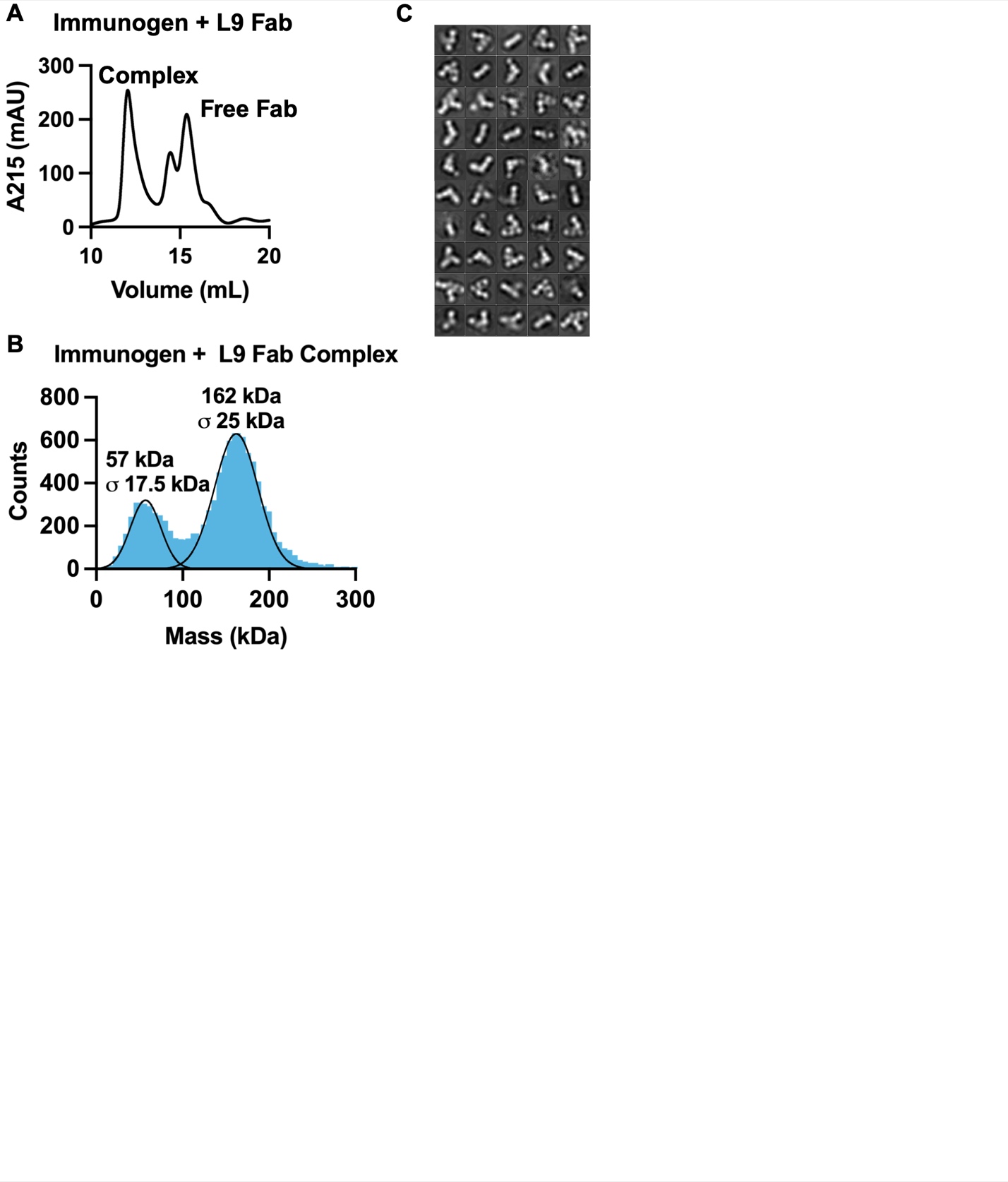

**Figure S6.** Purification of L9-M-TIM complex. (A) SEC trace of L9-M-TIM complex. (B) Single particle mass photometry of L9-M-TIM complex. (C) All 50 negative stain classes of L9-M-TIM complex.

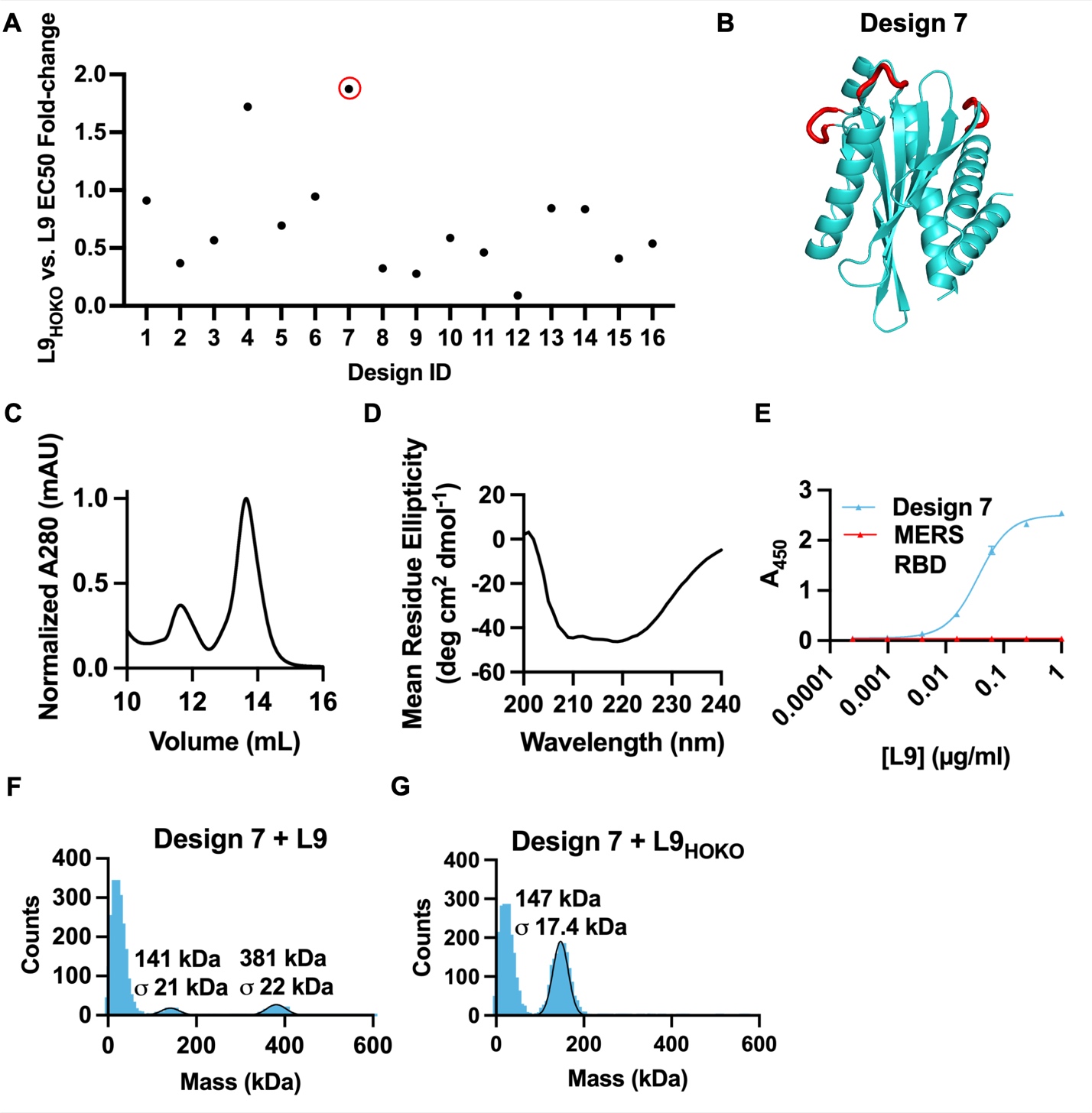

**Figure S7.** Biophysical characterization and antigenic profiling of 3-NPNVD design 7. (A) L9 vs L9_HOKO_ EC50 ratio of top 16 3-NPNVD designs. Red circle denotes design 7, the design with the largest change in binding to L9_HOKO_ vs L9. (B) AF2 prediction of design 7. Red residues represent the 3 NPNVD repeats. (C) SEC trace and (D) circular dichroism of design 7. (E) L9-design 7 ELISA binding curve. MERS-RBD was used as a negative control. (F) Single particle mass photometry of design 7 in complex with L9 and (G) L9_HOKO_.

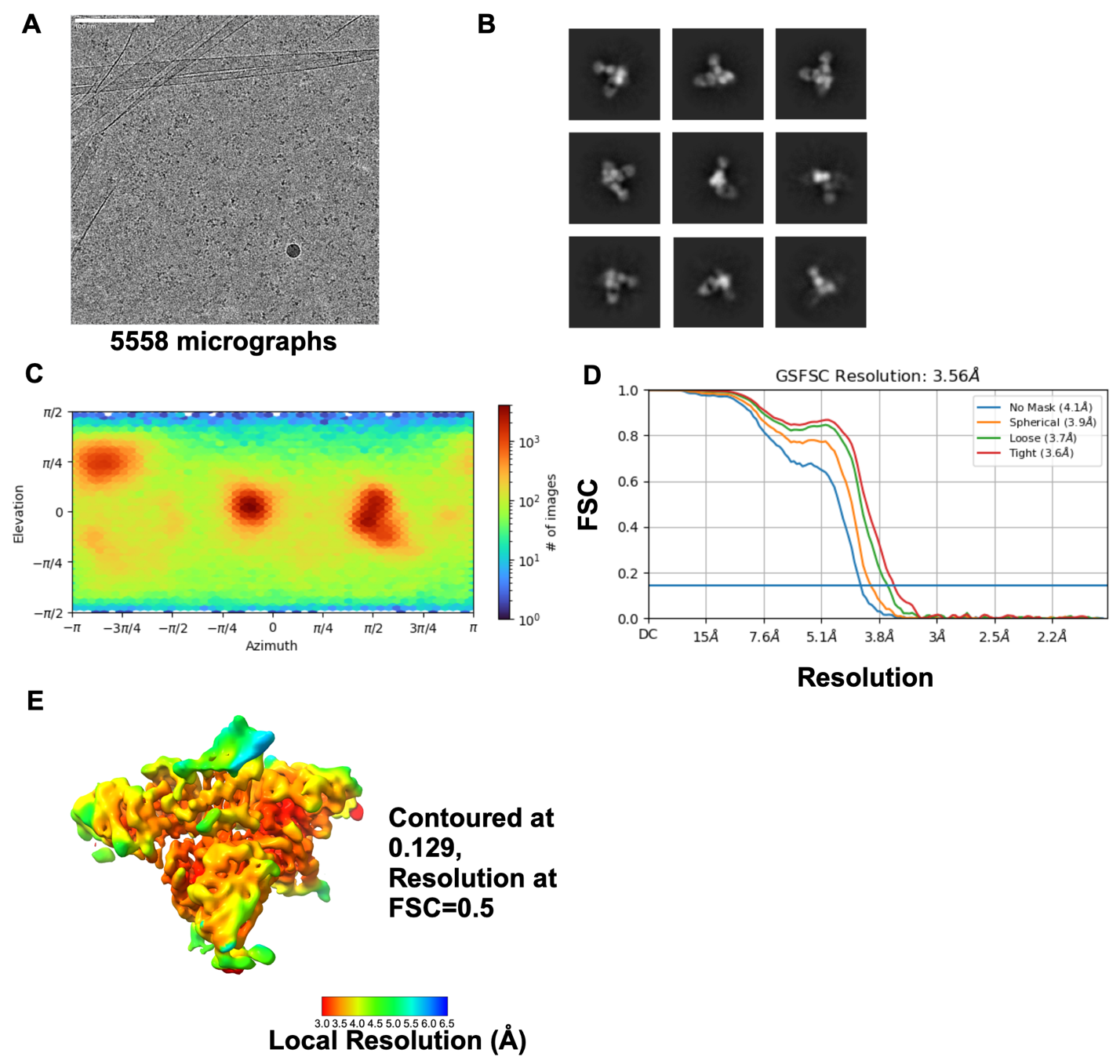

**Figure S8.** Cryo-EM details of L9-M-TIM structure. (A) Representative micrograph. Scale bar = 100 nm. (B) Representative 2D classes of L9-M-TIM complex used in 3D reconstruction. (C) Heatmap showing orientation of all particles used in final 3D reconstruction. (D) Gold standard Fourier shell correlation resulted in a 3.6 Å overall map. (E) Local resolution of the final unsharpened map is shown contoured at 0.129 with a FSC cutoff of 0.5

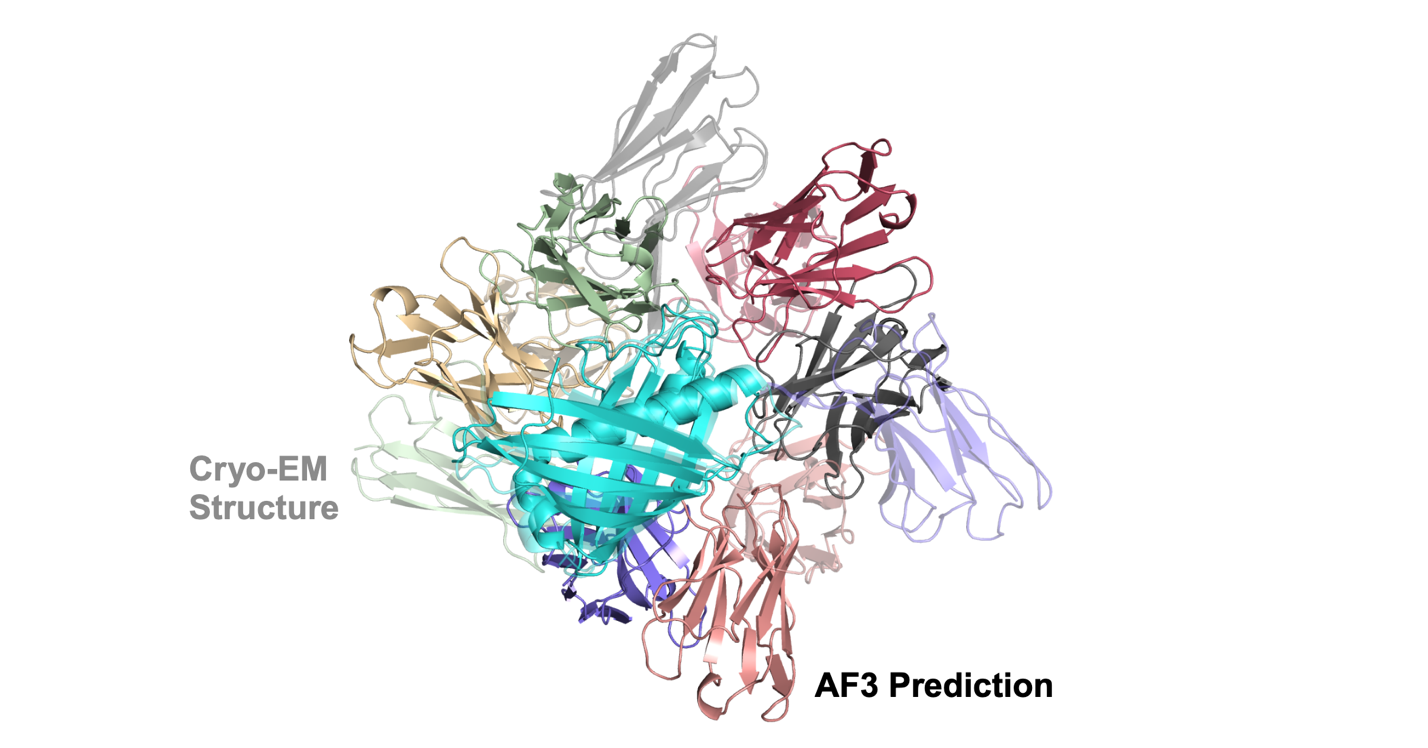

**Figure S9.** Comparison of L9-M-TIM cryo-EM structure vs AlphaFold3 prediction of the complex, aligned on the M-TIM structure. Cryo-EM structure is transparent while AF3 predicted structure is opaque.

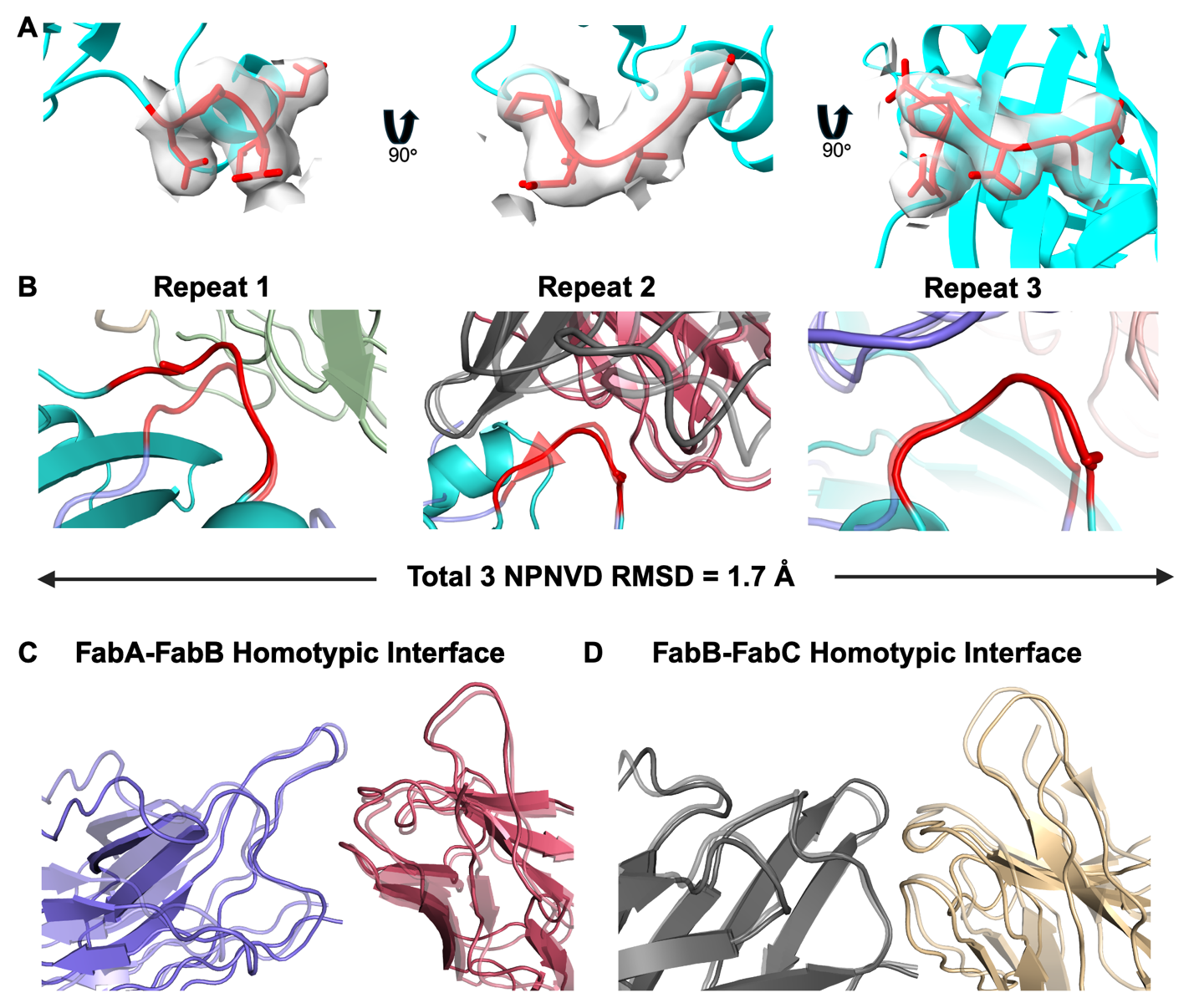

**Figure S10.** Structural characterization of important epitope-paratope and homotypic interactions. (A) Representative model fit in EMReady map density for minor repeat bound to heavy chain C and light chain D, three orthogonal views. (B) Overlay of each of the 3 NPNVD repeats from M-TIM with the 3 NPNVD repeats from CSP in PDB 8EK1. 8EK1 structure is transparent, while M-TIM structure is opaque. (C) Overlay of FabA-FabB and (D) FabB-FabC homotypic interactions between L9-CSP and L9-M-TIM structure. L9-CSP structure is transparent and L9-MTIM structure is opaque.

**Table S1.** Sequences of one minor repeat designs.

| Design ID | Sequence |
| --- | --- |
| 1 | MEIETITITDGSSVAEKCVELTKKYPGYVVYGINPNVYEVSEELIEDMAADVRAGLKVIIVGEDKERCEEVAEKVRKLAEGGHHHHHH |
| 2 | GFTPEEQLTIDYVNAKITVENYELLKEEAKKNPNVIVTKLDEVNYKAAKAFLKKYKVDEEKVKKKLEEILKGGHHHHHH |
| 3 | MEIEVVTEEEGKELLEKCRKEGKDIITLTINPNVSKEKAIKRIKEAIESCEADGSKLVITAPEELLPYAKELLEKAGGHHHHHH |
| 4 | DIKVTYKNGSCKEAYEDEAENYLSGGDYKKEYFCINPNVINISIAKEKVYEAYANGNTELVVYCPENLLPKVKKEIEEALAEIKKKGGHHHHHH |
| 5 | VETFEKTFKNEEEAVKFAKELFEEALKKGYTSITLESKGITIKVDLNPNVFDDSDLIEAVLAAYDDAKAAGVPLKVTLYKSGGHHHHHH |
| 6 | SSAKVIFVESDYNDVLLLTTKIIREEKVNYIKSYYINPNVILSDEEWLELAEQLLEDEKLAAELGGVVVIGVPDERTKEKLLEYIEKAKAGGHHHHHH |
| 7 | MVLSEEEIEELKKLIEEAREKGGTVIITLTNPETGEVVTYSYRVNPNVTNIEPVLEELKNIIEAGKLKGWNIKLELVVSPGGHHHHHH |
| 8 | MEIKIMTKEEVEKLAKEKGWKVYFSPPTSDFELALGEGVDKGADIVAITFNPNVLPDPGTLLNGLKRAKKLAEKYGIEFVIVSENPNELLGWLKSTGGHHHHHH |
| 9 | MDLSKLSEEEIKKLAKELYEKLYELSLKYGRLKITFYIGDYVETMTLNPNVPIRESTLERFEGLLLEIKYSGEDATLVIEAENNKEKITIVFKNGGHHHHHH |
| 10 | GPEIIGLVIEPGKHIDEEVIKKVNEIVRKLLKEGKKGIIVTQINPNVFGSAEKVAELAYETGETVVAVSDSEEELKKAKKLLEELLKKREEEKGGHHHHHH |
| 11 | GNFTLNVEKLKHPYEIKVTIKKKDGTTETTTSVANPNVRTDVDAVAKAAAEKVKAALDAGYDVTLTVGENMAKVVEKALKEAAGGHHHHHH |
| 12 | GPEIIEEWYPYSEKSKEKVKKILEEKGAKYGKVYYYNPNVFDQYVEIRLYKLENGKLYLISNRSEYYKRNGGHHHHHH |
| 13 | KKRILIVTTTDPEHKSKAEKKVKELRKKGIEAHLYFVNPNVFKPKSYFLEIFLAYDVDKVLVWKSLPEHHLKAIEEAAKELGIELEVIEGGHHHHHH |
| 14 | VKKVFKRTFYAEAETTEELKKKIKEAQEEAKKNGTVKSILLLRQPGNPNVVITEDLPIDMSDEEIENFCKELQEYEDLTGTKSKYKIYFVSEVPSGGHHHHHH |
| 15 | SSEKKVVLLILLVSMSDEEFKEYLENYKEEVDVLIKKYGVEKVVVIIASNPNVLISEEQKELLEKFVKELKEKGVEVIFAKNEIEAIKLYNKLLGGHHHHHH |
| 16 | LEKKLSLLNKLRQLANGTITGYDSGGRLVTIHLNPNVFYTDEEMEELLEKLDDLKGVPEEDKKRVEELKKVIKELLEIKKKEGGHHHHHH |
| 17 | KLKVVSIEVVKKEIKPSGSITLTVEVTYENGEKVLHTVTMYHNPNVETQEAEVKCLDCGCSIIDYLSNSDESFENAAFKLVLSCGGHHHHHH |
| 18 | MKVKLVLDEEALEEMLKKLGAKKEKVSPGLTKYTIEYTPDTKISLYVRRNPNVYPGAVYTEKIEATDEEVAKKFVKEELYNIARGGHHHHHH |
| 19 | KKKRAGIFSKDYPSTKEKNLEAYKKLSEELKKKGYGIVEYYSVNPNVSAESKAEKAYKSGYYDLVIVLENGKVKKTLKGGHHHHHH |
| 20 | MTFTVTFDTEEDGAKLLSLMEKLGAAKVLKTEVKNGKMTAVIEVNPNVFLTPAEVLEAGVSVEGAKINKELNERVRKELASSGGHHHHHH |
| 21 | MMVVRITITVYKKPKPEEIKAALERLQELLKPYGARVELHVEIGNPNVFTNTIIELSSGQVYTIYTHSSLSLEEKVKLVEPILEKAVKAASGGHHHHHH |
| 22 | PKINISETIENSKLPEYFEALLGQNISHITITVNPNVLAEPSQYPKILESYEKLEKYCKESGATLTLSEELKRLLERIREELEKKKGGHHHHHH |
| 23 | EKETVTIEMPVEELTVEKLVELLKKSGKITIKVLTESGKEITKEFTYNPNVDSDRVKVAEEALEWVKEYASKGAKVTIIIEVPKGGHHHHHH |
| 24 | SRTVHLQFDDVEGLKLVKAEVLRLIEAGVDKVLISKNPNVIPTPAEKKEYDEILKELLSLPKVELVEPPPGGHHHHHH |
| 25 | TTTLSINLTDDTVTVIIKALPGTNATLTYEQNGSKVTKVLKGNPNVPSTLEVTLNNAKGKTLTVSASGPGCVSVTLITPNKTIFLPDICGGHHHHHH |

**Table S2**. Sequences of three minor repeat designs.

| Design ID | Sequence |
| --- | --- |
| 1 | VVKIKFKVELKCEEITVKTPEKTFKLDVCEATVTICTSDGKKYKFKVKITRGERNPNVDPASLEAADKVIVVAAVLAYIKGLPSGSEVYVRAWATPNPNVDNIVELYIEENGEKYLIVSVMLNPASGQVSPANPNVDPETVAKGAGVSLIIDDWFKVEVESESSEEELEELKKLVKKYLEQGGHHHHHH |
| 2 | MEEINLEVEVKVLKHKNGRVSILVIVRSDNYAAYAEINPNVDLHCANTDPVCLEALVTLLETLLTTLVRLLEERGIDVEELEAILWWPDCNPNVDPIFRLVHNGSPFDLELQQELVETLQKKIKELNKKNPKLKYPPNVRLANPNVDDETELANTGRLVPITITVRKREEMEPEIEELLKRFEELEKEEKGGHHHHHH |
| 3 | MLTLFTKFTSKLVSVEPGKHKYSIYDKETGYTFDITLTKSYDPATGKTYYKFDIDPKHANPNVDLKPVTVTAEKEDRPPREREFLGRRLTTDSLRGTVSYNPNVDLRFVAVRARSEDGHRAYAVVLVDGAPREEEAEAALDALLLRIHKEGLWDSEEVLALVQVGNPNVDPSEWAFSPLMLETMGLAAAAAVIALAEVEELEEKAAGGHHHHHH |
| 4 | MVEIEKKKFSVEEENNKVTLTLCKEKTDGTVTCLTVEWEYEVNPNVDPNDQKAFKEITILSLKEMLGKNFEEFVKLNGKKLKVSIKATITSNPNVDNEVRLKVKSEDGKIFELCDMYRCDPDEPCIKTPRKCNPNVDLETIAKRTGSRVVEEEFTMNLKVEKKKEEIELEGGHHHHHH |
| 5 | ETETFKRELTTPDFKFSLEITIEKIDEFPNLFLINAEGVATPDNPNVDFPPIKFASSVIVVITENEDGVKEIQVNSVTALYIQANPNVDMMLIKRNIKLENLPKELQEKIIKLMEEKEKELVEKIKKWSGKKYVDAPNPNVDPSEYDPVVRECLKQLLLVQEELEEALNKLIEELKPEFIKRVGGHHHHHH |
| 6 | MKITSVYILIKCSEKLKPILEPIAKEIEREKGYKTEIIANPNVDESSWLTIAVLTIVEENGKISASISVRAKHVKNPNVDASLTTSLLPSSEELIKETKELLDEIWKRIAKALGLPTSVREANPNVDPEEIAKINPEMANGYREILCAICEMKELLPNLIKEMREEEAKGGHHHHHH |
| 7 | SAEALLAEAVDHLAAGDLNAAARTLVELIGERVRKENPNVDVGDTSFEELADGTARATLHCPPDLSVVVLIRREEAGGPTVVILALIVANPNVDLALVSVLVVRREDGRLRVYVASAKIENGKAKLFEPTDPELKAIVDKAVQDALKLYGVRLFNPNVDATTQCKNAGISTGKMEFLADVVDEVLQALAAALAARDGGHHHHHH |
| 8 | MERLEEFKKELEEKLEELEKELEELVKETIKETIEKIKSNPNVDTSNLKIEVSIESSREFLSYDEEKGTMKARITLYVKINITPNPNVDLAATITIEVEVESNEKTGEYKILKWTFSVEYSRGYNPNVDLKTWLKRNPSLYIAFLLLEEIEENLKKAISEFVEEKKKKGGHHHHHH |
| 9 | EKLVIVIGTQPGDKEKTERYKKLIREALKELGIEAEIYEIECNPNVDFETCVKILVKKALKYAAEKGAKAIAIGINTGVKEVLLPNPNVDITIEAGPEIIGDIVLIIINELEDGKVVIESYVLYSKYNPNVDNPTLSLIPRKIKTKETSNDDEALLATLKETIKESIEELEKLEEERKKLEGGHHHHHH |
| 10 | MSKEEIEKKKFMEELFEEAKKLMEELIKEINSNPNVDESLKMKLEEKVIETEENTYRFVLGKGEKVTIAVGMSPEGKICVHIQYNPNVDANICSSNEDLPFCEECIELLNKEFGLNIIPSFLNPNVDLETQAKNLGINVFLLEDAKRLHDALNKKKEELLEKLKKGGHHHHHH |
| 11 | AEEEIKKILDEIIKLLVEAGQPLIIDRSFMSPEILKYIEERAKKIPKAEIKDISELEFNPNVDYTVIEIRVSGLSAEETAAAREVTEKVVKHIQKVLKEKGSRAKVSINPNVDFFIPVSKARELLLEAVKKAAERGLPVIVSVVTTDRGRLLVVVIIEGADVAAEVVNPNVDADEILKLLGIAPWNIVIMTLFFHDPEAVAAAKGGHHHHHH |
| 12 | MKKIKIYIRFHLSNNPEDLKKDKENKEKIIKKLKEYLEKKGYTLNVEYISNPNVDFAPIYDVEETYEILKTIIDIFSSGNTIAISVASFDNPNVDCILTKSIDNYSVLMLLIPRKDGNSLLIVVVSKDENLIKKAYDYFVTEKGLPRCNPNVDLSGLANPYIRERMKLALAEAEELVELLKKLDGGHHHHHH |
| 13 | MCTVVIVIACEEEHLPLAEEIAERIRKELGLEVELFQCNPNVDLELFIDIAISEVIDKYNCDKIIVIVIVDESKTLSSLINPNVDNFSLACNSCVKAGCYTVDICLIAISREEVEPGVYRVDILCKTLICNPNVDDSKRKPYTLSQSFSSKNMTLEELLRYIKSLLGGHHHHHH |
| 14 | MKYAYFSGKGTREEVSDRIFELHLKAEKEGNKNFVALFLIEPNPNVDPNIIIVVLVTSNSDTVKVEAKFDGLEVKIEVKNPNEKGLTIVVIAFDLRNPNVDSGAYRAVIEDGKISFSGVTVNVEIIDEKTGVRIKGTRSTNPNVDLETWLKLGGSPTWVNLTRLAEAIARYIAESILRKLKDYKEKEGGHHHHHH |
| 15 | MIIEVELDLPGYTVKITLEQVSKDTYKVKIKITHNPNVDPIYKILIKTPGEEKEIDDKDSELEYTIKSDGEEVILEFKNKNPNVDGDVTIKVSFGKVEIIAKTDIGKYSTLKIENDKIDISLANPNVDVETLRKNGAYLQAHIKESLVNTYKELLKKKKEGGHHHHHH |
| 16 | MKKLKIHVLVAYNSRDTEKKPKLCALAERIAAFLKEKYPDAEVTVTEVENPNVDSTLVAELASLDLIDELKKESEETGNEITLVSVYFRSNPNVDGHTCEAACEAIKAGFRYIVVSANDQQGSKIRLKKIAPNPNVDDKTTTLIDSKSFDTLEDALAEAVEIIEKEPQLVVELMVGLGGHHHHHH |
| 17 | SLEELLKKKKKIEEEIKKEILKRIDEIAEQEAANPNVDPAVRDIIVASLEVIKELCKELSLEEIKYLKVRLIVDPNPNVDNLLFLSLENEKQKKMIKELEERTGLRFKLANPNVDFETYKKNGYNGLLALEDTVKEIAKEVLEEYEKKLGGHHHHHH |
| 18 | STIKKELKIEKTMGLDIVKLFIEDEESKTKTIIEVSINPNVDKKARDVMLKTAKELFEEWLKDDSLKEFEKILKKAITMALVMLWLARLALNPNVDLVLRVSLSPEVGHLIDIVLEAIEEAKELVAKRQEIKGRQPRLNPNVDCEDFELPEACAKLYVEKLDENTLLISVGSAGIEVDITIKKEKLGGHHHHHH |
| 19 | ELRLEILREVVKKTKIQITVSESLYEELMKANKEDLEAVLKMLGVELETNPNVDKATGINKLAIYFLELVNLVLDTLTEEEIPKVVKVLKIIAELVRKYANPNVDVLVHVTLNGKVTAVRIDQALELRAEALEKGEKVTMEFAYNPNVDIETIKKISTPEAALKHEFALEAKKIVSKIKYEVEKELEKLKKEGGHHHHHH |

**Table S3.** Cryo-EM single-particle data collection and refinement statistics.

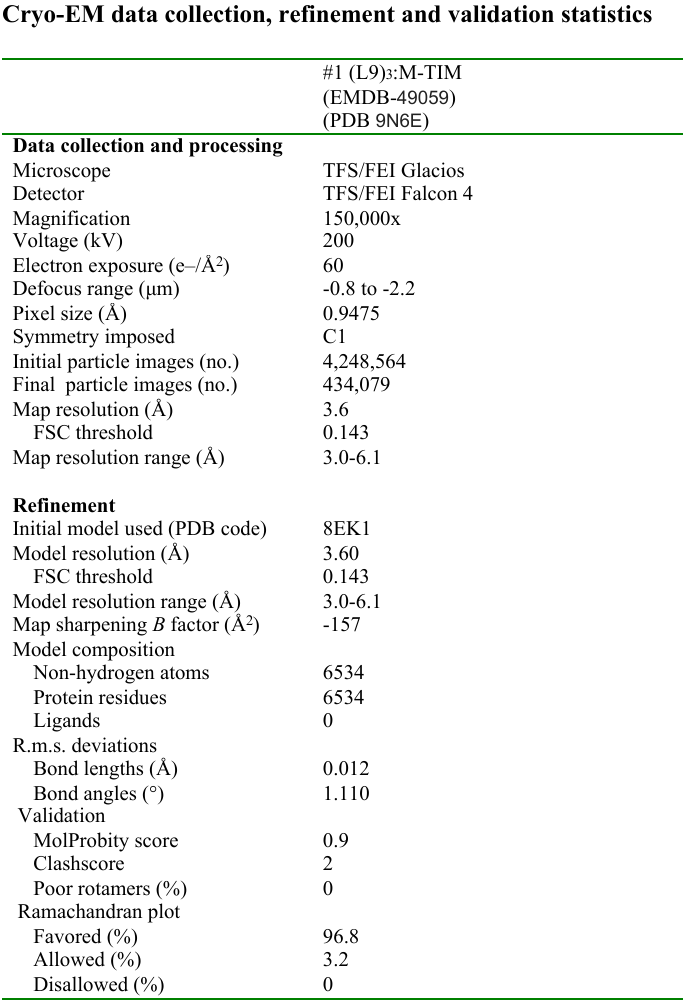
